# Sierra: discovery of differential transcript usage from polyA-captured single-cell RNA-seq data

**DOI:** 10.1101/867309

**Authors:** Ralph Patrick, David T. Humphreys, Vaibhao Janbandhu, Alicia Oshlack, Joshua W.K. Ho, Richard P. Harvey, Kitty K. Lo

## Abstract

High-throughput single-cell RNA-seq (scRNA-seq) is a powerful tool for studying gene expression in single cells. Most current scRNA-seq bioinformatics tools focus on analysing overall expression levels, largely ignoring alternative mRNA isoform expression. We present a computational pipeline, Sierra, that readily detects differential transcript usage from data generated by commonly used polyA-captured scRNA-seq technology. We validate Sierra by comparing cardiac scRNA-seq cell-types to bulk RNA-seq of matched populations, finding significant overlap in differential transcripts. Sierra detects differential transcript usage across human peripheral blood mononuclear cells and the Tabula Muris, and 3’UTR shortening in cardiac fibroblasts. Sierra is available at https://github.com/VCCRI/Sierra.

## Background

Regulation of cellular transcriptional activity includes changes in the total expression level of genes as well as alternative usage of gene architecture. Alternative, or differential, usage leads to expression of alternative mRNA transcript isoforms, which we refer to here as differential transcript usage (DTU). Forms of DTU can include alternative splicing (AS), such as through exon skipping or retained introns, alternative 5’ promoter usage or 3’ end use, or changes in 3’UTR length through the use of alternative polyadenylation (APA) sites. RNA sequencing (RNA-seq) studies have revealed high levels of alternative transcript usage between tissues, with 95% of multi-exon genes estimated to undergo AS among human tissues [1]. Similarly, APA is widespread among the mammalian genome, and is estimated to occur in most genes [2]. While the extensive use of AS and APA among tissues is documented, DTU between the diverse sub-tissue cell-types revealed in recent years by single-cell RNA-seq (scRNA-seq) is relatively unexplored, nor is the regulatory logic well understood.

There is a strong impetus to develop strategies for studying DTU at the single-cell level, given that studies to date have found the different forms of DTU to play significant functional roles across many biological contexts. Functional consequences of DTU can include changes to mRNA stability, localisation, protein translation and nuclear export as determined by AS [3] or APA in 3’UTRs [4]. DTU plays roles in differentiation and organ development through AS [5, 6], including intron retention (IR) [7, 8], a form of AS that appears widespread in mammals [9, 10]. DTU is also relevant in disease contexts. As examples, AS is linked to disease through miss-splicing caused by genomic variants [11] while 3’UTR shortening has been reported as widespread in cancer cells [12] and has been found to repress tumor-suppressor genes [13]. Alternative usage of introns is also linked to cancer both through intronic polyadenylation [14] and IR [15, 16]. Overall, studies to date reveal a pattern of widespread alternative mRNA transcript usage of different forms and functional consequences across tissues, development and disease contexts.

The advance of scRNA-seq technologies opens up new avenues for deeper exploration of DTU at the level of single cells [17]; however, there are technical features of the high-throughput nanodroplet technologies like the 10x Genomics Chromium platform, including low depth and limited gene coverage, that makes detecting DTU a non-trivial task. Some scRNA-seq protocols such as Smart-seq provide read coverage across the gene and therefore enable the analysis of alternative isoform expression [17]. For example, analysis of neural Smart-seq scRNA-seq data has demonstrated alternative splicing events at the single-cell level in the brain [18] and methods have been specifically developed for analysing alternative isoform expression in scRNA-seq experiments that have reads spanning the transcript [19–21]. The above studies utilise increased transcript read coverage at the expense of lower-throughput profiling of cells; however, high-throughput nanodroplet technologies like 10x Chromium have become the technology of choice for many scRNA-seq experiments due to the capacity to profile thousands of single-cell transcriptomes at low cost. Currently, gene-level expression data is primarily utilised in analysing 10x Chromium data; however, the enrichment of 3’ ends in barcoded and polyA-captured scRNA-seq means that APA and alternative 3’-end usage can be explored, with the potential to unveil additional levels of information among cell types currently masked when only considering an aggregate of each gene. Despite polyA-captured nanodroplet scRNA-seq experiments now routine, to the best of our knowledge there is no computational pipeline described that can leverage such datasets to identify DTU between cell types.

We present here a novel computational pipeline for unbiased identification of potential polyadenylation (polyA) sites in barcoded polyA-captured scRNA-seq experiments, and evaluation of DTU between cell populations. We demonstrate that we can identify DTU – which we define as any change in relative transcript usage, including differential exon usage, alternative 3’UTR usage and changes in 3’UTR length – between cell-types and across different tissues and disease contexts. We demonstrate the application of our pipeline using public data generated from a variety of 10x Chromium scRNA-seq experiments on human PBMCs, murine cardiac interstitial and enriched fibroblast cells from injured and uninjured hearts and a multi-tissue atlas from the Tabula Muris. We validate our approach by comparing DTU calls from cardiac scRNA-seq data to bulk ribo^−^ RNA-seq of matched cell populations derived from FACS and find a significant overlap in DTU genes from both the scRNA-seq and bulk RNA-seq experiments. Our analysis can detect multiple types of DTU among cardiac interstitial cells types, including alternative 3’ end usage, APA and even alternative 5’ start sites that are validated from the matched bulk RNA-seq samples. We further apply our method to detect 3’UTR shortening in proliferating cardiac fibroblasts from injured mouse hearts and show not only that we can detect 3’UTR shortening in proliferating cells as observed previously in other tissues, but can detect shortening in related activated populations that are not proliferating – a granularity that is not possible to observe in bulk RNA-seq studies. We also provide an *in vivo* validation of candidate 3’UTR shortening genes using real time quantitative RT-PCR. Finally, we apply our approach to 12 tissues from the Tabula Muris, presenting an initial atlas of cell type-specific DTU across mouse tissues. Our analysis pipeline is implemented as an open source R package, named Sierra, available at https://github.com/VCCRI/Sierra.

## Results

The Sierra R package contains a start-to-end pipeline for identification of used polyadenylated sites in scRNA-seq data, differential usage analysis and visualization (Methods). Briefly, the Sierra pipeline starts with a BAM file, such as that produced by the 10x Genomics CellRanger software, and the reference GTF file used for mapping (Figure 1). Based on the observation that aligned reads from 10x Chromium scRNA-seq experiments fall into Gaussian-like distributions, peak calling is run to identify local regions with high read coverage, or ‘gene peaks’, within genes that correspond, for example, to different potential polyA sites or other transcript features. The peak coordinates are utilised to construct a new reference file of genomic regions, enabling a unique molecular identifier (UMI) matrix of peak coordinates to be built for a supplied list of cell barcodes. Each of the gene peaks is annotated according to the genomic feature it falls on (3’UTR, exon, intron or 5’UTR) and proximity to sequence features including A-rich regions and the canonical polyA motif – enabling discrimination of likely cases of internal priming to A-rich sequences from true polyA sites.

**Figure 1.**
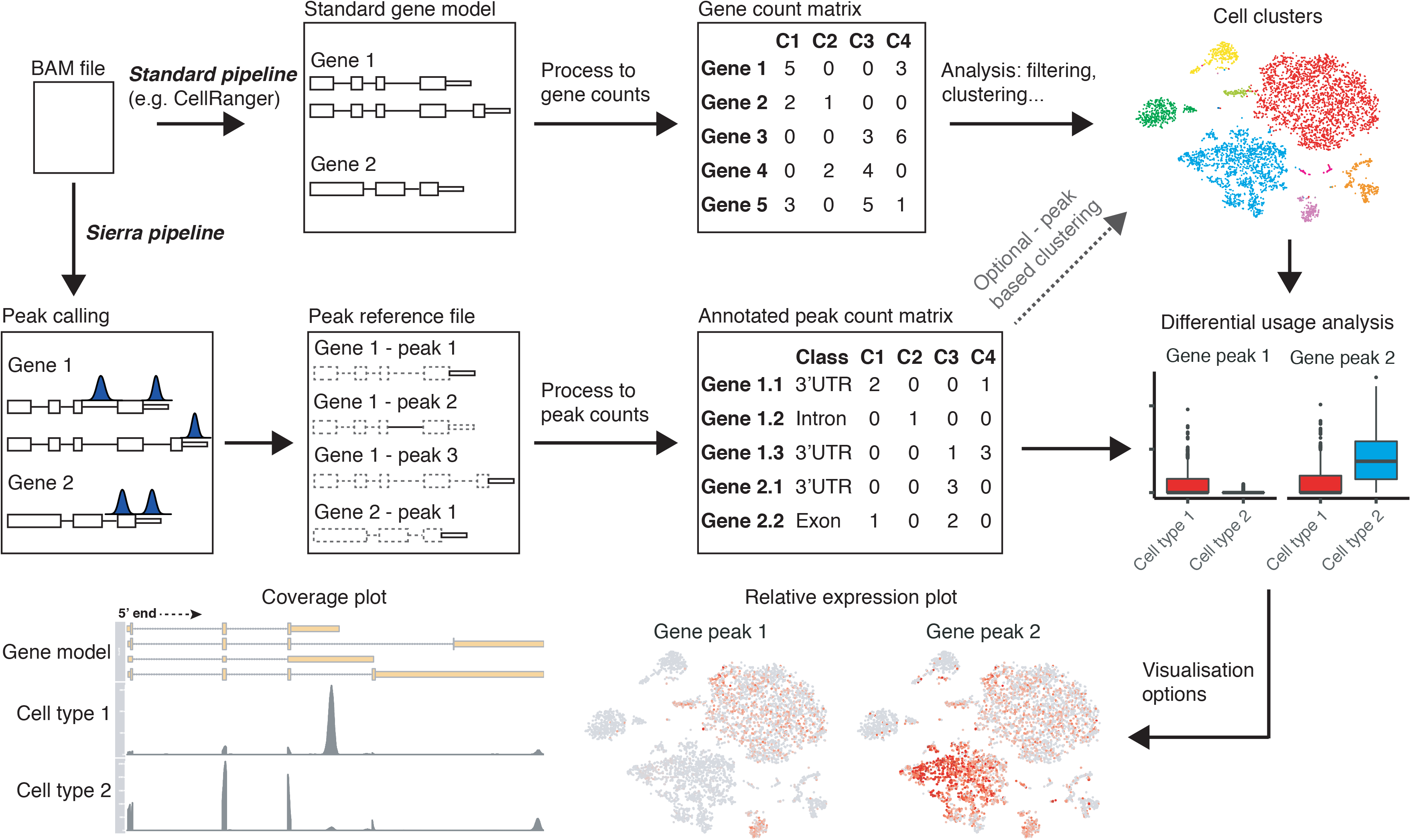
Sierra workflow. Sierra starts with a BAM file produced by an alignment program such as CellRanger. Standard gene-level work-flow (top row) involves using a gene model to produce a matrix of gene-level counts used for clustering. The Sierra pipeline performs peak calling to identify coordinates corresponding to potential polyadenylation sites. Peak coordinates are used to build an annotated UMI count matrix for each gene peak. This new data can be used to identify genes showing differential peak usage, with visualisation options for plotting relative peak expression and read coverage across selected cell populations.

For downstream analysis, Sierra can utilise either the Bioconductor [22] SingleCellExperiment class [23] or the widely-used Seurat R package [24] to create objects containing the peak counts and annotation information required for DTU testing, in addition to pre-determined cell identities/clusters and dimensionality reduction coordinates (t-SNE or UMAP) for visualisation purposes. For Seurat users, Sierra can directly import cluster identities and t-SNE/UMAP coordinates over from a pre-existing Seurat object, allowing for straight-forward analysis of pre-defined cell types.

To test for DTU between cell groups we utilise the differential exon usage method developed for bulk RNA-seq, DEXSeq [25], but applied to peak coordinates, whereby a gene will be called as DTU if it shows significant change in the relative usage of peaks. Cells within each cell group are first aggregated into a small number of pseudo-bulk profiles, which are used as replicates in DEXSeq, enabling computational efficiency and statistical power for DTU identification. Sierra also contains several functions for visualisation of peak expression from DTU genes (Figure 1). These include the plotting of relative expression between two or more gene peaks, where the expression of a set of gene peaks is transformed according to their relative usage within each cell population (Figure 1; Methods). Relative expression can then be plotted on, for example, t-SNE or UMAP coordinates as with gene expression data. Finally, in order to aid in interpretation of DTU between cell types, we provide functionality for gene-level plotting of read coverage across defined single-cell populations. Single-cell populations (e.g. clusters) are first extracted from an aggregate BAM file according to cell barcode, generating population-level BAMs. Read coverage for DTU genes can then be visualised at a single cell-population level (Figure 1). The coverage plots, which also show a gene model, allow for deeper interpretation of the nature of the DTU detected.

### Features of Sierra data

We applied Sierra to 19 publicly available datasets representing a variety of common scRNA-seq experimental settings: two human PBMC datasets, (a 7k cell and 4k cell) from 10x Genomics (https://www.10xgenomics.com/), total non-cardiomyocyte cells (total interstitial population [TIP]) from uninjured (sham) hearts or hearts at 3- or 7-days following myocardial infarction (MI) surgery [26], enriched cardiac fibroblast lineage cells (*Pdgfra*-GFP^+^) also from MI and sham mouse hearts [26] and the Tabula Muris [27], with twelve tissues from Tabula Muris analysed in this study (Table S1).

Sierra typically detected 30,000 – 50,000 peaks covering 10,000 - 15,000 genes across the datasets tested (Table S1). Most genes had a small number of peaks, typically with a median of 2 peaks per gene (Table S1). In the PBMC 7k dataset for example, over 4,000 genes were called with 1 peak, followed by ∼2,000 and ∼1,000 genes with 2 and 3 peaks, respectively, while a minority of genes were called with 20 to 100+ peaks (Figure 2A). Considering the genomic features associated with these peaks, we found that for genes with 1-2 peaks, the majority of these fell in 3’UTRs (Figure 2B), while genes with larger numbers of peaks tended to show more intronic peaks. For genes with over 20 peaks, on average ∼75% of these were intronic (Figure 2B). These metrics were consistent in both the PBMC 4k and TIP datasets (Figure S1A-D). We next stratified the peaks according to genomic feature type, and examined how increasing stringency of cellular detection rates (i.e. only considering peaks expressed in some *x*% of cells) affected the feature-type composition of peaks. With no filtering, we found that the largest number of called peaks were intronic, followed by 3’UTRs (≥ 0% detection rate; Figure 2C and Figure S1E,F). Progressively stringent filtering of peaks according to cell detection rates showed that intronic peaks tended to be detected in a smaller number of cells (Figure 2C and Figure S1E,F). The substantial presence of intronic peaks is in agreement with previous observations made about RNA molecules containing intronic sequence in 10x Genomics Chromium data [28], and likely corresponds to pre-spliced mRNA.

We compared the expression characteristics of the peaks with gene-level expression data from CellRanger (Figure S2A-D) and found a strong correlation between gene expression and expression of peaks in 3’UTRs as expected, with weaker correlations in intronic peaks for both 7k PBMCs (Figure S2A) and the cardiac TIP dataset (Figure S2C). We also compared gene and peak expression using mean expression versus dispersion plots, calculated with Monocle [29]. We noticed a wider range of dispersion values in peaks compared to genes for both datasets, although intronic peaks partially explain this, with a higher dispersion range among more lowly expressed genes (Figure S2B,D). Finally, we annotated each peak according to whether it was proximal to an A-rich region or the canonical polyA motif (Table S1). We found 3’UTR peaks had the highest percentage of proximity to the polyA motif (on average 47%), while 5’UTRs had the lowest (average of 5%). Intronic and exonic peaks also had low levels of polyA motif proximity (average of 9% and 10%, respectively). Conversely, 3’UTR peaks had the lowest proximity to A-rich regions (average of 10%), while intronic peaks had the highest (50%), with exonic and 5’UTR peaks showing an average of 28% and 18%, respectively (Table S1).

**Figure 2.**
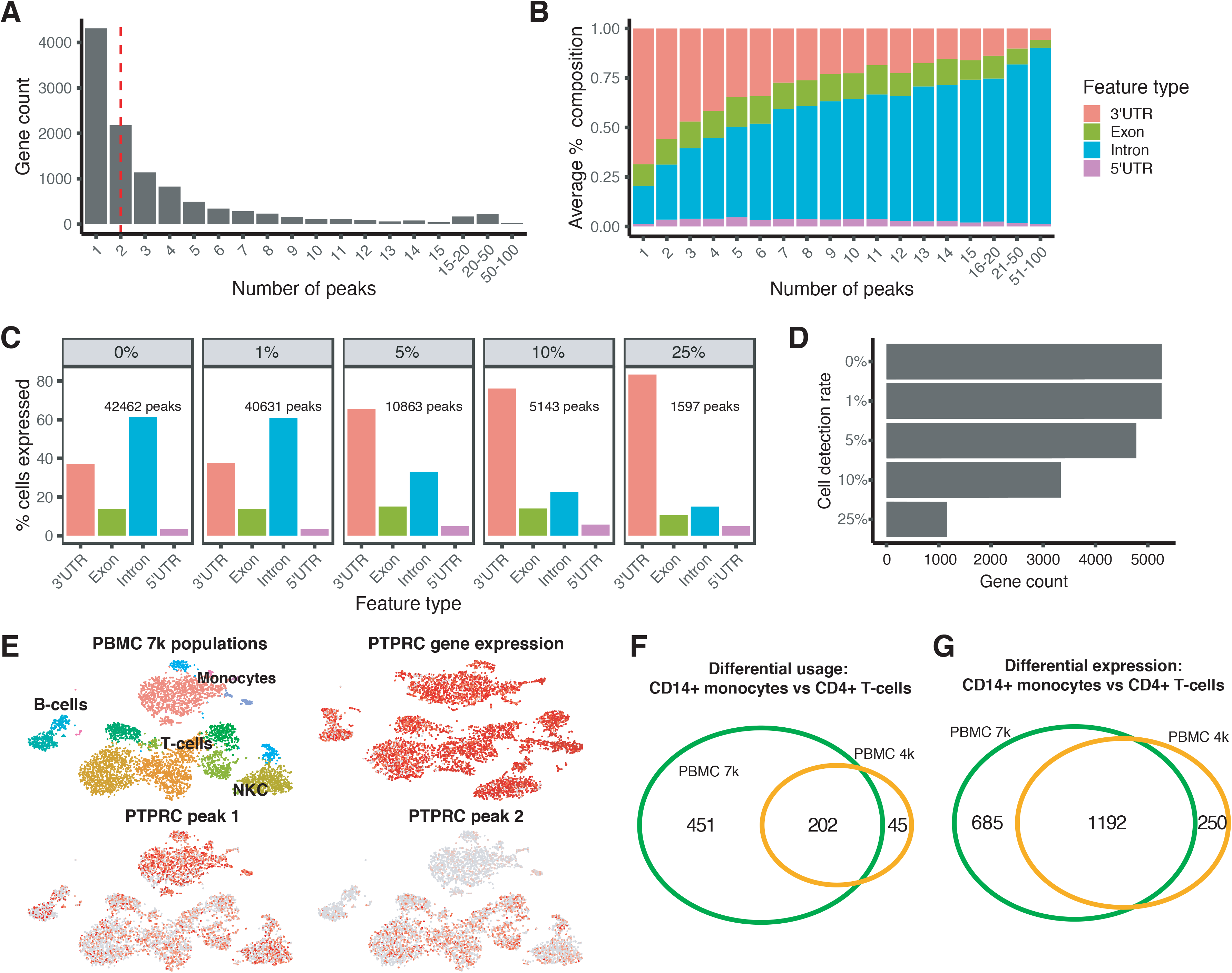
Representative feature of Sierra data from a 7k cells PBMC data-set. (A) Counts of genes according to number of detected peaks. Dotted red line indicates median number of peaks. (B) Average composition of genomic feature types that peaks fall on, according to number of peaks per-gene. (C) Percentage of cells expressing each genomic feature type with increasing stringency of cellular detection rates for peaks. (D) Number of genes expressing multiple (≥ 2) 3’UTR or exonic peaks with increasing stringency of cellular detection rates. (E) Comparison of *PTPRC* gene expression across cell populations on t-SNE coordinates with peaks identified as DU in monocytes. (F,G) Overlapping genes from a CD14^+^ monocyte vs CD4^+^ T-cell comparisons for the PBMC 7k and PBMC 4k data-sets for (F) DTU genes and (G) DE genes, visualised with [60].

### Differential transcript usage among human PBMCs

We next considered the extent to which we could call DTU between human PBMC cell populations as defined by gene-level clustering. Seurat clustering of the 7k PBMCs yielded 16 clusters including several populations of monocytes (Mo), CD4^+^ T-cells, CD8^+^ T-cells, B-cells, two natural killer cell (NKC) populations and several minor populations. As DTU testing only applies to genes with multiple peaks, we evaluated how many genes had multiple 3’UTR/exonic peaks expressed across the PBMC 7k cell populations at increasing cell detection rates. When filtering for peaks with expression in at least 10% of cells, there were over 3000 genes with multiple peaks. This dropped to just over 1000 genes when requiring peaks to be detected in 25% of cells (Figure 2D). For the below analyses we used a 10% detection-rate cutoff. We applied DEXSeq [25] to call peaks exhibiting differential usage (DU) between cell types after aggregating cells within cell types into a small number of pseudo-bulk profiles to create pseudo-replicates (Methods). We restricted our testing to peaks falling on 3’UTRs or exons.

We performed DU analyses both between clusters and aggregated groups of cells (e.g. all CD4^+^ T-cells vs monocytes). We readily detected significant DTU genes using our pseudo-bulk replicates with DEXSeq between cell types (*p_adj_* < 0.01; *LFC* > 0.5). We detected the largest numbers of DTU genes when comparing cell groups from different lineages; for example, Mo against lymphoid populations including B-cells (BC), T-cells (TC) or NKCs (see Table 1 for representative examples). When comparing populations Mo1 and CD4^+^ TC1, 825 distinct peaks were called by DEXSeq as DU, representing 492 DTU genes (i.e. a gene containing at least one DU peak is classified as a DTU gene). In contrast, DEXSeq detected 83 DU peaks corresponding to 63 DTU genes when comparing populations NKC1 to CD4^+^ TC1 (Table 1). Of the DU peaks, on average 20% were near an A-rich region, and potentially due to internal priming, while an average of 32% flanked the canonical polyA motif. Among the DTU genes, there were known examples of alternatively spliced genes in immune cells, including *IKZF1* (Ikaros) and *PTPRC* (CD45) [30]. Although *PTPRC* gene expression is ubiquitous among immune cell types, we found we could distinguish clear patterns of alternative peak usage in monocytes compared to other cell types (Figure 2E) according to t-SNE visualisations of relative peak expression, demonstrating that we are able to detect cell type gene expression activity masked when only considering an aggregate of the reads across a gene. Plotting read coverage for *PTPRC* and *IKZF1* revealed examples of DU peaks corresponding to alternative 3’-end use between the Mo and TC populations (Figure S3A,B).

**Table 1.**
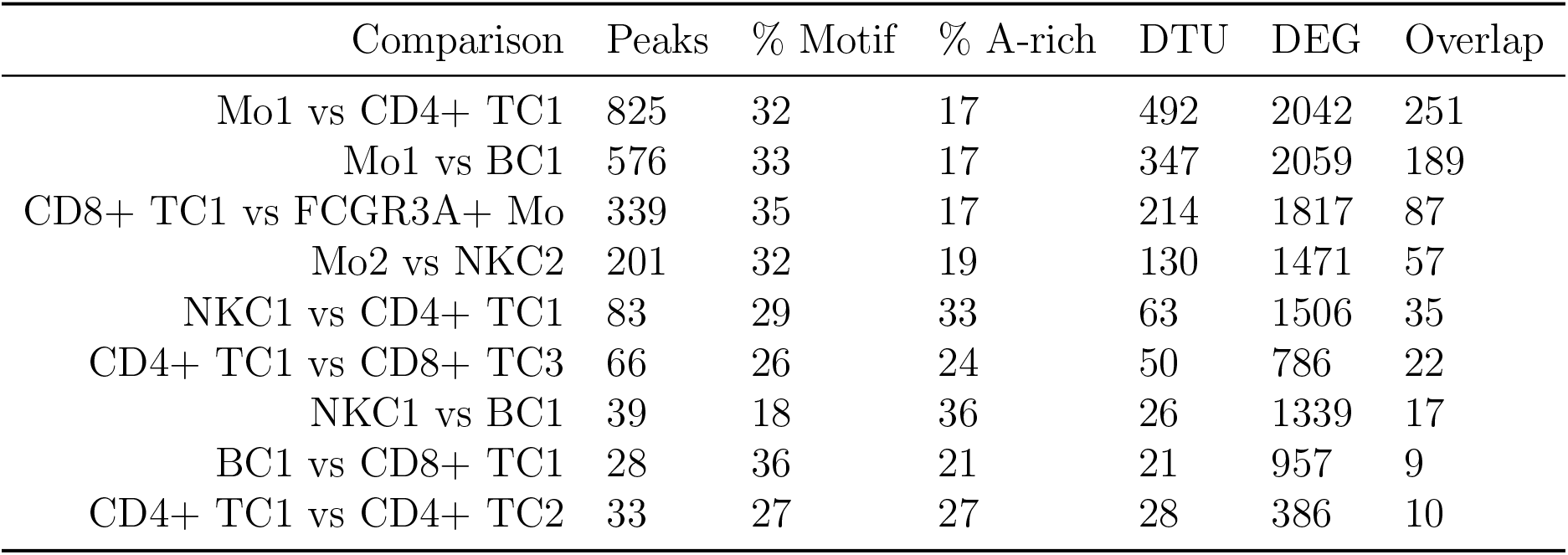
Differential transcript usage (DTU) examples on the PBMC 7k data-set. Shown are the specific cluster comparisons performed, number of peaks called as differentially used (DU), the % identified as proximal to the canonical polyadenylation motif, the % proximal to an A-rich region, the number of unique called DTU genes, the number of genes called as differentially expressed (DEG), and the number of overlapping genes from the DTU and DEG lists.

We next explored whether a relationship existed between DTU genes and differentially expressed (DE) genes using counts derived from CellRanger. We compared DTU genes to DE genes from the same population comparisons using the Seurat *FindMarkers* program with MAST testing [31]. For the PBMC 7k dataset, we found on average that 50% of the DTU genes overlapped with DE genes. We also found a strong positive correlation between the number of DTU and DE genes (Spearman’s correlation test; *ρ* = 0.79; *p* = 3.2*e* − 51). Finally, we evaluated the reproducibility of calling DTU genes by comparing the PBMC 7k DTU genes with genes identified by the same cell-type comparisons in the 4k PBMC dataset. For example, comparing CD14^+^ Mo to CD4^+^ TCs, there were 653 DTU genes in the PBMC 7k dataset and 247 in the lower-depth PBMC 4k dataset, with 202 overlapping (Figure 2F), representing 82% of the DTU genes from the PBMC 4k analysis. For comparison, we performed the same analysis with DE genes and found a similar level of overlap (Figure 2G). Across all comparisons, we found that on average 60% of DTU genes were matched between PBMC 4k and PBMC 7k datasets, while for DE genes an average of 80% were matched (Table S2). Thus, the majority of DTU genes can be found in a replicate experiment, and at levels not far below DE testing, indicating that our method of detecting DTU genes is reproducible.

### Patterns of differential transcript usage in the mouse heart

We validated Sierra DTU calls by comparing single-cell populations from the mouse heart [26] to a bulk RNA-seq dataset of matched cardiac populations isolated with FACS [32]. Importantly, the bulk RNA-seq experiment used a ribo^−^ protocol instead of polyA^+^, which makes this a unique validation resource that does not have the 3’ bias of the scRNA-seq. The cardiac scRNA-seq experiment was performed on the total non-cardiomyocyte compartment of the heart (total interstitial population [TIP]) and reported several interstitial cells types including fibroblasts, endothelial cells (ECs) and numerous sub-populations of leukocytes [26] (Figure 3A). In addition to cardiomyocytes, the bulk RNA-seq dataset contains sorted ECs (CD31^+^), fibroblasts (CD90^+^) and leukocytes (CD45^+^). These populations were isolated from adult mouse hearts at 3 days post-sham or myocardial infarction (MI) surgery; the TIP scRNA-seq experiment contains a sham, and two MI time-points: MI-day 3 and MI-day 7. Both datasets therefore contain comparable populations and conditions (sham and MI-day 3). To compare with the scRNA-seq we first applied DEXSeq to calculate DTU genes between an aggregate of the scRNA-seq fibroblast cells, ECs and leukocytes from the sham hearts (i.e. pairwise comparisons between these cell lineages) and the sham populations against MI leukocytes (Table S3). We did not include MI fibroblasts or ECs as these populations are diluted in the scRNA-seq due to an overwhelming influx of monocytes and macrophages at MI-day 3 [26].

**Figure 3.**
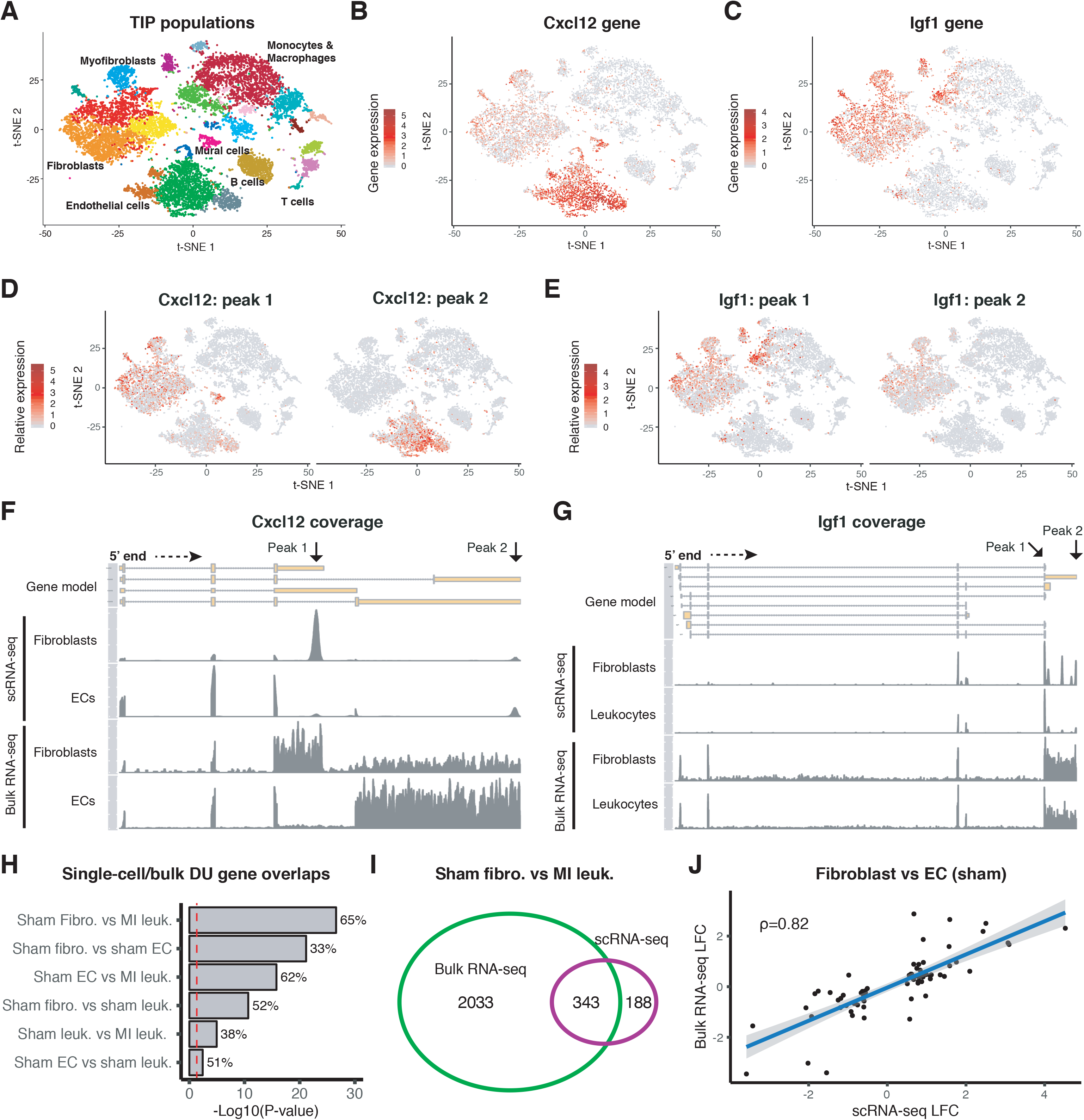
Comparison of differential transcript usage between cardiac scRNA-seq and bulk RNA-seq populations. (A) t-SNE plot of the cardiac TIP cell lineages. (B,C) Gene expression visualised on t-SNE for (B) *Cxcl12* and (C) *Igf1*. (D,E) Relative peak expression visualised on t-SNE for example DU peaks between (D) sham fibroblasts and sham ECs: *Cxcl12*; and (E) sham fibroblasts and MI leukocytes: *Igf1*. (F,G) Read coverage plots across the *Cxcl12* and *Igf1* genes for (F) single-cell and bulk fibroblast and EC populations (*Cxcl12*) and (G) single-cell and bulk sham fibroblast and MI leukocyte populations (*Igf1*). (H) Fisher’s exact tests on the number of overlapping DTU genes detected from scRNA-seq and bulk for different cell-type/condition comparisons. Shown are the −log10 p-values and the percent of single-cell DTU genes overlapping the bulk. Red line indicates the significance (0.05) threshold. (I) Overlapping genes between single-cell and bulk RNA-seq from the sham fibroblast and MI leukocyte comparison. (J) Log fold-change comparisons for DU peaks identified in both the single-cell and bulk RNA-seq for the sham fibroblast vs EC analysis.

We found clear examples of DTU masked when considering an aggregate of the gene. As examples, top DTU genes from fibroblast to EC and fibroblast to MI leukocyte comparisons were *Cxcl12* and *Igf1*, respectively. *Cxcl12* was observed to be expressed in fibroblasts, ECs and mural cells (Figure 3B), while *Igf1* was expressed in fibroblasts and macrophage populations (Figure 3C). When we compared relative expression of DU peaks, we found we could distinguish clear patterns of specificity between fibroblasts and ECs for *Cxcl12* (Figure 3D) and between fibroblasts and macrophages for *Igf1* (Figure 3E). We plotted read coverage across *Cxcl12* for the fibroblast and EC populations and compared them to the bulk RNA-seq of sorted fibroblasts and ECs. This showed that the peak upregulated in fibroblasts corresponded to a 3’UTR of a short transcript isoform of *Cxcl12*, while the top EC peak corresponded to a longer isoform (Figure 3F). The difference in transcript isoform expression between fibroblasts and ECs was also observed in the bulk RNA-seq, confirming our observations made in the single-cell data (Figure 3F). The fibroblast and EC transcripts correspond to the *Cxcl12*-α and *Cxcl12*-γ isoforms, respectively, with their protein products showing alternative localisation and functional properties (see Discussion). We were able to experimentally validate *Cxcl12* transcript isoform expression by specifically targeting the different 3’ exons using real time quantitative RT-PCR (qRT-PCR) (Figure S4A,B).

In the case of *Igf1*, the two DU peaks corresponded to proximal and distal sites on the same 3’UTR, relative to the terminating exon, and are annotated in current RefSeq gene models. The peak distal to the terminating exon (peak 2) was preferentially expressed in fibroblasts, and from the bulk RNA-seq we could also observe that fibroblasts preferentially expressed this longer 3’UTR in *Igf1* (Figure 3G), demonstrating that we can detect APA corresponding to changes in 3’UTR length. We could observe other examples of APA from DTU genes between fibroblasts and leukocytes, including *Tfpi* and *Tm9sf3* (Figure S5A,B). We also detected examples of alternative 5’ start sites in *Lsp1* and *Plek* (Figure S5C,D), demonstrating that Sierra detects DTU corresponding to a variety of alternative transcript expression events.

We next asked how many DTU genes detected from the single-cell comparisons could also be detected in the bulk RNA-seq. We used the peak coordinates mapped to 3’UTRs and exons as a reference to generate counts from the bulk RNA-seq and again applied DEXSeq to determine DTU between the same cell-types and conditions as with the scRNA-seq data. For all comparisons we found that there was an overlap between the scRNA-seq and bulk RNA-seq DTU genes greater than expected by chance (Figure 3H; Fisher’s exact test; *p* < 0.05), after using the set of genes with multiple 3’UTR/exon peaks expressed in the relevant scRNA-seq populations as a random-expectation background. The comparison with the largest and most significant overlap was between sham fibroblasts and MI leukocytes, with 343 out of 531 comparable DTU genes (65%) from the scRNA-seq analysis also observed as DU in the bulk RNA-seq (Figure 3H,I). We next compared the fold-change direction of the peaks called as DU in both the scRNA-seq and bulk RNA-seq experiments. For all comparisons we found a significant positive correlation in fold-change (Spearman’s correlation test; *p* < 0.05). The strongest correlation was found in the fibroblast vs EC comparison (Figure 3J; *ρ* = 0.82). We also considered whether filtering out peaks annotated as A-rich prior to the DTU analysis would improve the correlation with the bulk RNA-seq. We recalculated DU peaks from the single-cell populations, first filtering out peaks proximal to A-rich regions. For 5/6 of the comparisons there was no major change in the metrics used for the comparisons; however, for the sham EC vs sham leukocyte comparison we noticed the overlap increased from 51% (with Fisher’s exact test *p* = 0.004) to 56% (*p* = 0.001) and the Spearman correlation coefficient increased from 0.31 (with Spearman’s correlation test *p* = 0.005) to 0.4 (*p* = 9 × 10^−4^). Together, these results show that Sierra can detect multiple types of alternative mRNA isoform usage with a significant number of these corroborated by an independent bulk RNA-seq experiment.

### 3’UTR shortening in activated and proliferating cardiac fibroblasts

Proliferating cells have previously been observed to have, on average, shortened 3’UTRs [33,34]. We sought to determine whether we could apply Sierra to infer 3’UTR shortening at the single-cell level. In the heart, fibroblasts become activated and proliferate following MI, with the peak of proliferation occurring within days 2 to 4 post-MI [35]. scRNA-seq of enriched (*Pdgfra*-GFP^+^) murine cardiac fibroblasts at day 3 post-sham or MI has revealed several sub-types of cardiac fibroblasts [26]. In the uninjured heart, predominant fibroblast populations can be distinguished on the basis of the expression of *Ly6a* (Sca1) – referred to as Fibroblast Sca1-low (F-SL) and Sca1-high (F-SH) (Figure 4A) [26, 36]. Following MI, there is the expansion of a pool of activated fibroblasts (F-Act) leading to a population of actively cycling fibroblasts (F-Cyc) (Figure 4B). In between F-Act and F-Cyc in pseudo-time is an intermediary activated population, F-CI (cycling intermediate), that does not express markers of actively proliferating cells.

**Figure 4.**
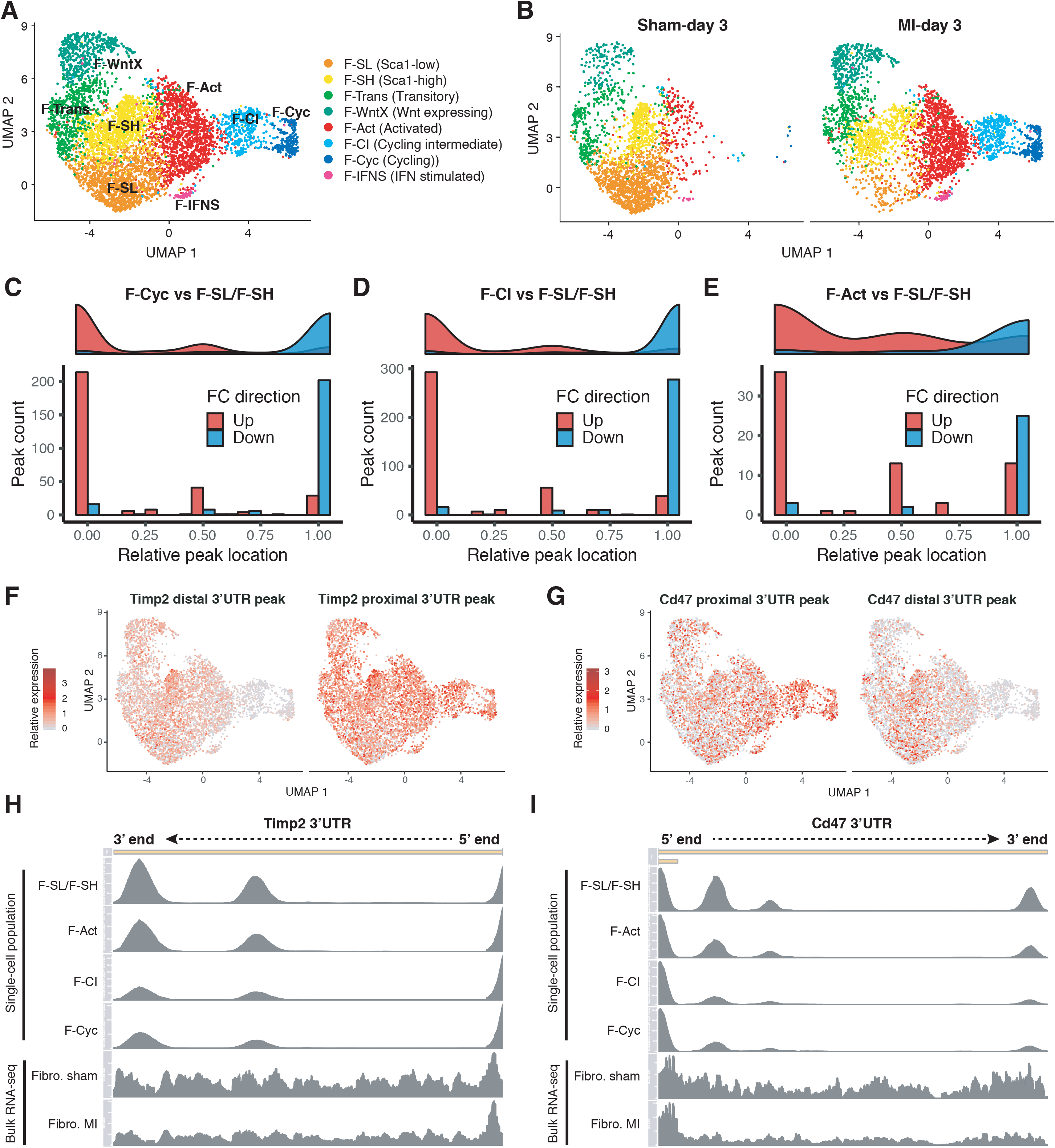
3’UTR shortening in activated and proliferating cardiac fibroblasts following MI. (A,B) UMAP visualisation of fibroblast populations from *Pdgfra*-GFP^+^/CD31^−^ mouse cardiac cells at 3 days post-sham or MI surgery showing (A) an aggregate of all cells and (B) the UMAP plot separated according to condition. (C-E) Counts of 3’UTR peaks showing differential usage according to their relative location to the terminating exon. Location of 0 indicates the peak most proximal to the terminating exon, with 1 representing the most distal. Comparisons performed are for (C) F-Cyc against F-SL and F-SH combined, (D) F-CI against F-SL and F-SH combined and (E) F-Act against F-SL and F-SH combined. (F,G) Relative expression of peaks most distal and proximal (to terminating exon) for (F) *Timp2* and (G) *Cd47* as visualised on UMAP coordinates. (H,I) Read coverage across 3’UTR for select single-cell fibroblast populations from sham (F-SL/F-SH combined) and MI (F-Act, F-CI, F-Cyc) data-sets compared to bulk RNA-seq of FACS-sorted fibroblasts from sham and MI conditions for (H) *Timp2* and (I) *Cd47*.

We investigated whether we could infer 3’UTR shortening in the F-Cyc population compared to the main resting populations, F-SL and F-SH. We applied DEXSeq to find examples of DU peaks falling on 3’UTRs, filtering out peaks tagged as near A-rich regions prior to DU testing in order to enrich for real polyA sites (Table S4). For the DU 3’UTR peaks, we considered all expressed and non A-rich peaks that occurred on the same 3’UTR and ranked them according to their relative location to the terminating exon. Each peak was given a score between 0 (most proximal to the terminating exon) and 1 (most distal), and we determined if there was a difference in relative location for upregulated and downregulated peaks. We reasoned that an increased number of upregulated peaks proximal to the terminating exon should imply a preference of shortened 3’UTRs, as well as a downregulation of distal peaks.

Comparing F-Cyc to F-SL/F-SH, we found 598 DU 3’UTR peaks representing 424 DTU genes (*LFC* > 0.5; *p_adj_* < 0.01). Comparing their relative peak locations, we found a strong shift towards proximal peak upregulation in F-Cyc, and a corresponding propensity for distal peaks to be downregulated (Figure 4C; Wilcoxon Rank-sum test; *p* = 1.1 × 10^−68^. We next considered whether the remaining activated populations also showed evidence of 3’UTR shortening (Table S4). Interestingly, we found that the intermediate F-CI population, which we have previously interpreted to be a pre-proliferative state showing strong upregulation of translational machinery compared to F-Act [26], had an even larger number of DU 3’UTR peaks (785; 545 DTU genes) than F-Cyc and also showed a strong pattern of proximal peak upregulation (Figure 4D; Wilcoxon Rank-sum test; *p* = 6.5*e* − 98). We directly compared F-Cyc to F-CI but found only 8 3’UTR peaks showing DU, suggesting the 3’UTR shortening is occurring in the F-CI population prior to cell cycle entry. We also evaluated F-Act relative to F-SL/F-SH and found a smaller number of DU peaks (106; 94 DTU genes), which showed a significant, albeit reduced, pattern of proximal peak upregulation (Figure 4E; Wilcoxon Rank-sum test; *p* = 5.5 × 10^−8^).

We compared the genes implicated in 3’UTR shortening to the bulk RNA-seq of sorted fibroblasts from sham-day 3 and MI-day 3 hearts described above [32]. While the proliferating cells would have comprised a minority within the bulk RNA-seq, the most significant examples of 3’UTR shortening should still be detectable in the bulk RNA-seq. Two of the top significant DTU genes were *Timp2* and *Cd47*, which showed clear patterns of relative higher proximal peak usage in F-Cyc compared to resting fibroblasts (Figure 4F,G). We analysed read distributions in the 3’UTRs, including isolated single-cell populations F-SH/F-SL combined (from sham hearts) and the MI populations (F-Act, F-CI and F-Cyc) and compared these to the bulk RNA-seq of sorted fibroblasts from sham and MI-day 3 hearts (Figure 4H,I). In both the single-cell MI populations, as well as the MI bulk RNA-seq, we observed a decrease in coverage in the distal regions of the 3’UTR for both *Timp2* (Figure 4H) and *Cd47* (Figure 4I). We tested for DTU genes from the bulk RNA-seq using peaks from the GFP^+^ fibroblast lineage cells compared to sham cells, restricting our testing to 3’UTR and non-A-rich peaks as above. We found a significant overlap in DTU genes with the F-Cyc (Fisher’s exact test; *p* = 1.6 × 10^−15^) and F-CI (*p* = 7.6 × 10^−15^) populations, but not for F-Act (*p* = 0.078). Comparing fold-changes, we found a highly significant positive correlation between the bulk RNA-seq, and the F-Cyc and F-CI populations (Figure S6A,B; Spearman’s correlation test; *p* < 1 × 10^−100^). Despite the smaller overlap in DTU with F-Act, the peaks that did overlap nonetheless exhibited a positive correlation (Figure S6C; *p* = 10^−4^).

### *In vivo* validation of candidate genes for altered 3’UTR usage

We performed an *in vivo* validation of candidate genes associated with changes in 3’UTR length in proliferating cardiac fibroblasts, as detected by Sierra, using qRT-PCR. We first sought to determine the levels of cardiac fibroblast proliferation in murine hearts at different anatomical locations (zones) following MI injury (Figure 5A), in order to define proliferative and non-proliferative fractions for qRT-PCR analysis. We utilized a mouse model (*Pdgfra*^tdTom^) that expresses a red fluorescent protein (tdTomato) in PDGFRα+ cells, allowing for quantification of actively cycling fibroblasts using imaging (Methods). We subjected 8-week-old *Pdgfra*^tdTom^ mice to MI and measured proliferation seen as incorporation of the nucleotide analog 5-ethynyl-2’-deoxyuridine (EdU) into the newly synthesized DNA in replicating cells (Figure S7A). We found EdU incorporation in lineage-traced (tdTomato^+^) cells was exclusively restricted to the infarct zone (IZ), with 12.07±1.76% of tdTomato^+^ cells also EdU^+^, and absent from sham hearts and the remote zone (RZ) in MI mice (Figure 5B). Thus, both the sham and RZ form non-proliferative controls, with the caveat that the RZ likely contains activated fibroblasts [26].

**Figure 5.**
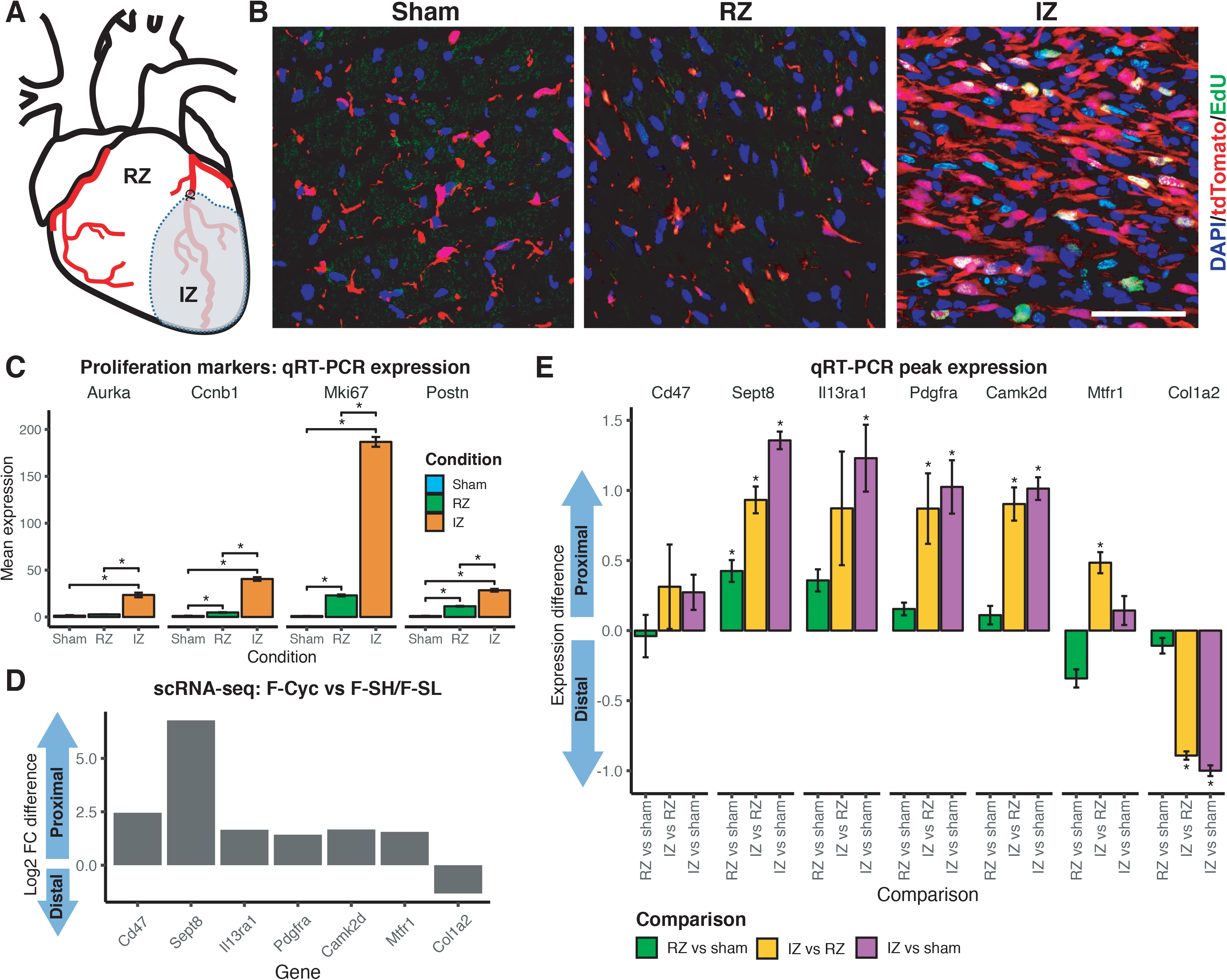
*In vivo* qRT-PCR validation of candidate genes with altered 3’UTR length in proliferating cardiac fibroblasts. (A) Diagram showing different anatomical locations of an MI heart: remote zone (RZ) and infarct zone (IZ). (B) Representative immunofluorescence images showing EdU^+^ cells in sham or indicated anatomical location of MI hearts. Scale bar indicates 50µm. (C) qRT-PCR expression of proliferation marker genes in *Pdgfra*-GFP^+^ cells sorted from sham hearts, RZ and IZ samples. Shown is mean expression and standard error (n=3), with stars indicating significant difference between comparisons (1-tail t-test; *p* < 0.05). (D) Sierra candidate genes exhibiting a shift to proximal or distal peak usage from scRNA-seq population F-Cyc in comparison to F-SH/F-SL. Shown is the difference in proximal to distal peak fold-change (log2). (E) qRT-PCR comparison of proximal to distal (P-D) peak expression from candidate genes in *Pdgfra*-GFP^+^cells (Figure S7) sorted from sham hearts, RZ and IZ samples. Y-axis represents ∆ ∆ (P-D) expression (log2; Methods) of sample comparisons RZ vs sham, IZ vs RZ and IZ vs sham. Shown is mean expression difference and standard error (n=3), with stars indicating a significant difference for the comparison (1-tail t-test; *p* < 0.05).

Furthermore, we performed MI surgeries on *Pdgfra*^+/GFP^ mice to validate the presence of altered 3’UTR usage as identified by Sierra. Given the zone-specific distribution of proliferating cells, we FACS-sorted GFP^+^/CD31^−^ fibroblast lineage cells from sham hearts, RZ and IZ of MI hearts at day 3 post-surgery (Figure S7B) and performed qRT-PCR assays. To validate that fibroblasts from the IZ were proliferative, we first confirmed that the expression of proliferation-associated genes (*Aurka*, *Ccnb1* and *Mki67*), as well as an activation marker (*Postn*), were increased in the IZ samples (Figure 5C). We tested differential 3’UTR shortening in 7 top candidate genes drawn from the Sierra analysis of proliferating (F-Cyc) vs resting (F-SH and F-SL) fibroblasts (Figure 5D) by designing PCR primers to cover the proximal and distal peaks on the 3’UTRs (Methods; Table S5). We selected 6 genes showing 3’UTR shortening (*Cd47*, *Sept8*, *Il13ra1*, *Pdgfra*, *Camk2d* and *Mtfr1*) and 1 that exhibited lengthening (*Col1a2*).

We further validated the peak expression, first by evaluating the difference (∆*ct*) in proximal versus distal expression within each sample, then comparing the ∆*ct* between samples to test for changes in proximal/distal peak usage (Methods). Comparing the RZ to sham samples, only one gene, *Sept8*, showed a significant difference (Figure 5E; t-test; *p* < 0.05). Given our findings above that activated fibroblasts (F-Act) exhibit some changes in 3’UTR usage compared to resting (sham) fibroblasts (F-SH/F-SL), observing that 1/7 genes show a significant difference between RZ and sham was not unexpected, and *Sept8* was in fact one of the DU genes detected in F-Act vs sham in scRNAseq comparisons (Table S4). However, we found the largest differences between IZ and RZ samples, and between IZ and sham samples, with 6/7 genes exhibiting significant (*p* < 0.05) shifts towards proximal or distal usage in line with the Sierra output (Figure 5E). The only gene that did not show a significant difference was *Cd47*, which was unexpected as we observed a clear shift towards proximal peak usage from single-cell and bulk RNA-seq of MI fibroblasts (Figure 4I). Nonetheless, the direction of difference seen was in line with our expectation from Sierra. Overall, these results provide strong experimental support for the DTU genes detected by Sierra.

### Clustering using peak-level expression

Our approach can clearly identify patterns of DTU when analysing single-cell populations defined through gene-level clustering. We next asked whether clustering on peak-level expression data could yield new information. We performed clustering using Seurat on the gene counts, with increasing cluster granularity through modification of the ‘resolution’ parameter in the Seurat *FindClusters* program, and compared the results to clustering performed on the peak counts, again selecting for peaks falling on 3’UTRs or exons (Methods).

We compared the clustering consistency between gene and peak counts using the Fowlkes and Mallow’s (FM) index and the Adjusted Rand Index (ARI). We found in general that the lower clustering resolutions, with fewer clusters, yielded higher consistency as determined by FM index and ARI (Table 2). We also noticed that the effect on cluster numbers returned from peak-level clustering tended to be dataset dependent. For the TIP dataset, there were fewer clusters relative to gene-level clustering (e.g. at a resolution of 0.6, there were 21 vs 25 clusters for peak-level and gene-level clustering, respectively). For the GFP^+^ dataset, there were consistently two additional clusters when using peaks, and for the PBMC 7k dataset, the numbers were the same (Table 2). Overall, using peak-level expression in place of gene-level expression does not appear to have a major impact on clustering results, particularly at lower clustering resolutions.

**Table 2.** Comparisons for clustering results based on peak counts relative to gene counts. Columns from left to right show the data-set tested, the resolution parameter (*res*) for the Seurat *FindClusters* program, number of clusters returned from gene-level clustering, number of clusters from peak-level clustering, Fowlkes and Mallow’s (FM) index and Adjusted Rand Index (ARI).

| Data-set | Resolution | Clusters (gene) | Clusters (peaks) | FM index | ARI |
| --- | --- | --- | --- | --- | --- |
| TIP | 0.6 | 25 | 21 | 0.92 | 0.91 |
| TIP | 0.8 | 29 | 25 | 0.86 | 0.85 |
| TIP | 1.0 | 30 | 27 | 0.79 | 0.76 |
| PBMC 7k | 0.6 | 17 | 16 | 0.84 | 0.82 |
| PBMC 7k | 0.8 | 18 | 18 | 0.82 | 0.80 |
| PBMC 7k | 1.0 | 20 | 20 | 0.81 | 0.79 |
| GFP<sup>+</sup> | 0.6 | 10 | 12 | 0.75 | 0.70 |
| GFP<sup>+</sup> | 0.8 | 12 | 14 | 0.75 | 0.71 |
| GFP<sup>+</sup> | 1.0 | 14 | 16 | 0.66 | 0.62 |

We visually compared the clusters returned by peak and gene-level clustering by imposing the clusters on the same t-SNE coordinates calculated on gene expression in the TIP dataset (Figure S8A-D). We found broad consensus between the clusters returned for most of the cell populations, further indicating that in general, the use of peak expression was not leading to the identification of many different populations. Of the few differences, we noticed that at resolution 0.6, the peak-level clustering identified a small sub-population of ECs (cluster ‘17’; Figure S8B) that was not present in the gene-level clustering (Figure S8A). We calculated DTU between cluster ‘17’ and the main EC population (cluster ‘1’) and found only a small number of DU peaks (16) between these clusters; however, differential gene expression testing between clusters ‘17’ and ‘1’ showed that cluster ‘17’ corresponded to a minor population of lymphatic ECs upregulating *Vwf*, *Lyve1* and *Prox1* (Figure S8E). The presence of these lymphatic ECs was observed in the original analysis by marker gene expression [26], but as a subset of cells within a larger cluster. Thus, clustering using peak expression may allow for a finer-resolution identification of some cell-types.

### A tissue atlas of cell type-specific differential transcript usage

We applied Sierra to the Tabula Muris [27], a compendium of scRNA-seq datasets across mouse tissues, to construct an initial tissue atlas of cell type-specific DTU. Applying Sierra’s peak calling to each of the 12 tissues, followed by merging of peak coordinates, we obtained a total of 107,425 peaks across the whole dataset. To determine DTU across tissues, we considered tissues that contained at least two cell types with at least 100 cells, leaving 10 tissues for our analysis. We calculated DTU within each tissue by performing pairwise comparisons between each of the cell types. Stratifying the DU peaks across tissues for each of the cell types, we found the greatest amount of DTU in mammary gland tissue, with most cell types exhibiting over 2000 examples of DU peaks. The smallest amount was observed in the heart tissue, which only contained fibroblasts and ECs after filtering for cell number, and between which there were 106 detected DU peaks (Figure 6A).

**Figure 6.**
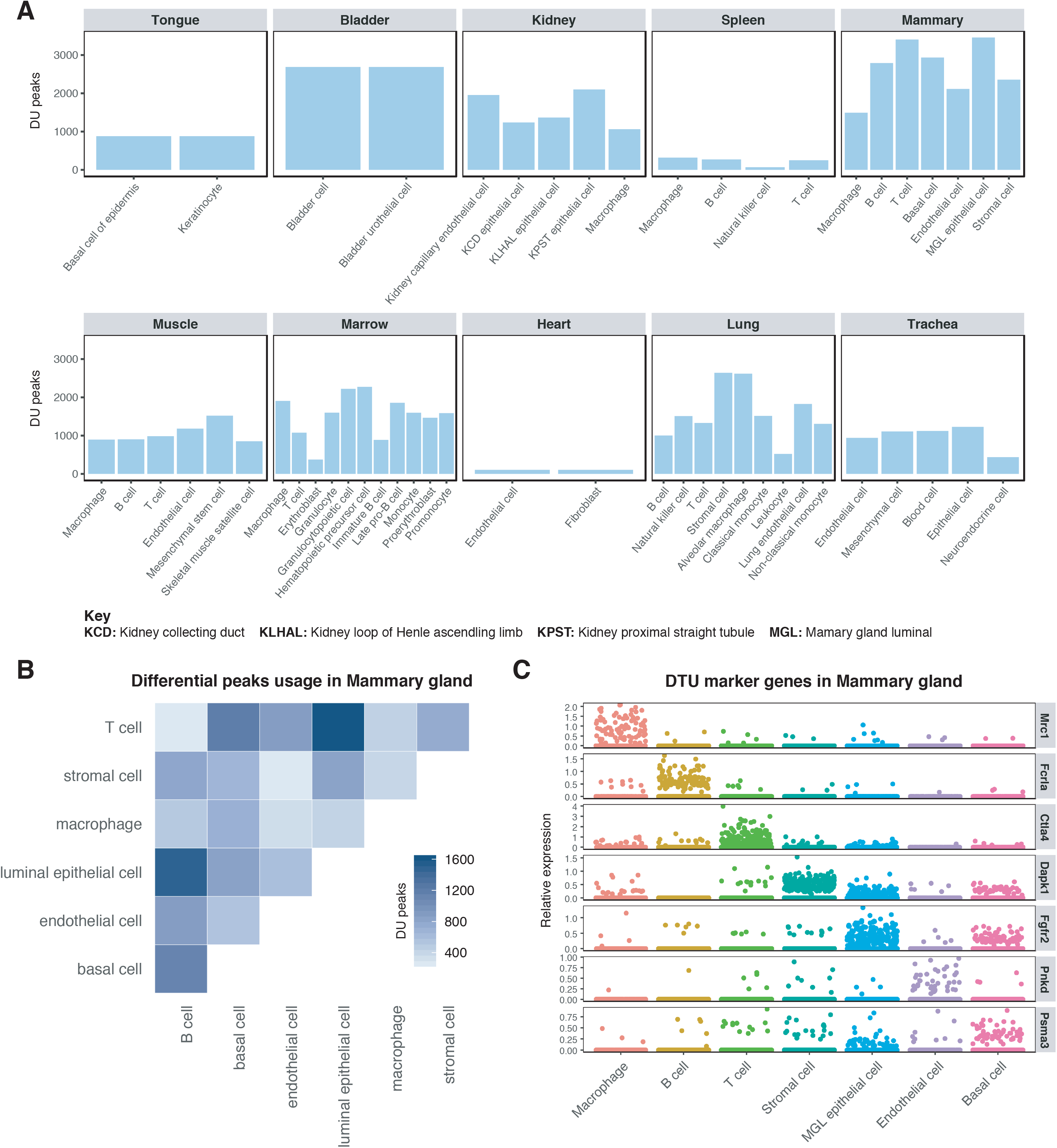
Detecting differential transcript usage across the Tabula Muris data-set. (A) Comparison of the number of DU peaks across cell types within each tissue. Only cell types with more than 100 cells are included in the analysis. (B) and (C) Mammary gland tissue results. (B) Number of DU peaks between cell types. (C) Relative expression plot of DTU genes between a cell type and all remaining cell types in the tissue.

As mammary gland tissue contained the greatest number of DTU genes, we focused more on this tissue type (Figure 6B,C). Across the mammary gland cell types, we found the largest number of DTU genes were called when comparing luminal epithelial cells to T-cells and B-cells. The smallest number of DTU genes was detected in the T-cell vs B-cell comparison. We next considered whether the Sierra DTU calls could be used to define “marker peaks” in an analogous manner to marker genes for cell types. Here we define marker peaks as peaks that are DU between the cell-type of interest and all other cell types. Applying marker DU testing to cell types from the mammary gland tissue, we defined DTU genes with high cell-type relative peak expression, including *Mrc1* in macrophages, *Ctla4* in T-cells and *Dapk1* in stromal cells (Figure 6C). We have made the processed Tabula Muris data available online (see Declarations).

## Discussion and conclusion

We have presented Sierra, a computational pipeline for discovery, analysis and visualisation of differential transcript usage in scRNA-seq data. Sierra is applicable to polyA-captured scRNA-seq experimental data such as those produced using the 10x Genomics Chromium system. Our method for detecting genomic regions corresponding to potential polyA sites enables a data-driven approach to detecting novel DTU events, such as alternative 3’ usage and APA between single-cell populations. We first determine the location of potential polyA sites by applying splice-aware peak calling to a scRNA-seq dataset, followed by annotation of the identified peaks and peak-coordinate UMI counting across individual cells. Finally, we use a statistical testing approach to determine genes exhibiting DTU with a novel pseudo-bulk approach to define replicates. While the software and analysis presented here is focused on the analysis of scRNA-seq data from UMI technologies, there is potential for our methodologies to be used in other enrichment based datasets such as single-cell ChIP-seq or ATAC-seq.

Using the Sierra approach we find thousands of genes that display significant DTU in publicly available datasets from human and mouse. Importantly, we provide several validations of our approach. Firstly, by use of qRT-PCR, we experimentally confirm select candidate DTU genes detected by Sierra. Secondly, we compare DTU calls made from scRNA-seq populations in the heart to DTU calls from ribo^−^ bulk RNA-seq of matched cardiac populations obtained with FACS. Both the qRT-PCR results and the significant overlap in DTU genes with the bulk RNA-seq points to the DTU detected by Sierra as representing real biology, and not technical artifacts of the scRNA-seq. For the bulk RNA-seq comparison, despite the significance of the overlap there were uniquely detected DTU genes in either the bulk RNA-seq or the scRNA-seq. These differences could be due to several reasons, both technical and biological. It is to be expected that more DTU genes will be detected in bulk RNA-seq data due to the increased depth and transcript coverage; however, there could also be examples of DTU genes that can more easily be detected with 3’-end-based scRNA-seq than with bulk RNA-seq, for example due to the transcript averaging effects in the latter, indicating a unique benefit of using 3’-end-based scRNA-seq for detecting some forms of alternative isoform expression.

The ability of Sierra to detect DTU provides the ability to contextualise prior known isoform information. For example, *Cxcl12* transcript isoforms are uniquely regulated and have known activities (reviewed in [37]). The *Cxcl12* isoforms detected by Sierra as DU between cardiac fibroblasts and endothelial cells correspond to α and γ isoforms, respectively. In the heart, the terminating amino acid sequence of Cxcl12-γ encodes a nuclear localisation sequence (NLS) and accumulates in the nucleolus, with any competing 5’ signal peptide being bypassed through the use of a non-canonical CUG start codon [38]. In other tissues, such as the brain, Cxcl12-γ is secreted, presumably because the 5’ signal peptide takes precedence over the 3’ NLS. Interestingly Cxcl12-γ is retained at cell surface due to its highly charged amino acid tail, which confers a 10x higher affinity for glycosaminoglycans when compared to other Cxcl12 isoforms [39, 40]. Cxcl12-α is the most widely expressed isoform but also is likely to have the shortest half life due to the 3’ end being a preferred proteolytic target of CD26 [37, 41]. A unique feature of Cxcl12-β is that the 3’ UTR sequence contains at least two bindings sites for miR-141/200 which have been shown to regulate only this isoform [42, 43]. The unique properties of Cxcl12 isoforms demonstrates the kinds of additional biology that can be inferred from scRNA-seq data when analysed by Sierra.

The flexibility of Sierra means that diverse questions can be asked about DTU. While we focus on mature transcripts in this manuscripts, the presence of intronic peaks means that questions about pre-spliced mRNA can be explored as well. The RNA Velocity approach utilises ratios of spliced and unspliced reads from scRNA-seq to estimate rates and direction of cell differentiation [28]. With Sierra, it should be possible to test for genes that show differences in relative usage of intronic peaks, as an indicator of changes in the expression-level of pre-spliced mRNA transcripts. Such analyses could be useful in differentiation contexts in identifying what genes are tending to be newly transcribed. There are additional applications that could be used for intronic peaks. One form of AS is intron retention, which has been found to have a role in cancer [15]. Intronic polyadenylation has also been linked to cancer through the inactivation of tumour suppressor genes [14]. The majority of intronic peaks from our analysis are annotated as proximal to A-rich regions, indicating that most will be due to internal priming, but by filtering for peaks more likely to represent true polyA sites, analysis of intronic polyadenylation represents a potential application of Sierra.

In conclusion, we have developed a novel computational pipeline for detecting differential transcript usage from 3’-end-based scRNA-seq experiments. Our novel approach to analysing scRNA-seq yields biological insights unobserved when considering only an aggregate of genes and allows new questions to be asked about the nature of transcriptional regulation between cells.

## Methods

### Data-sets

The locations of datasets used in this study are listed under Declarations. Previously generated bulk RNA-seq datasets [32] were downloaded, trimmed using Trimmomatic [44] and aligned using the two-pass STAR alignment method [45].

### Sierra pipeline

The Sierra pipeline is implemented as an R package and can be divided into the following main steps, each described in detail below. 1) Splice-aware peak calling is applied to a BAM file to identify peak coordinates. 2) If multiple datasets are being analysed, peak coordinates from multiple BAM files are merged together into one set of unified peak coordinates. 3) UMI counting is performed against the set of unified peak coordinates for a provided set of cell barcodes. 4) The peak coordinates are annotated according to the genomic features they fall on and, optionally, according to proximal sequence features corresponding to A-rich regions, T-rich regions or the presence of a canonical polyA motif. 5) Differential transcript usage analysis by applying DEXSeq to pseudo-bulk profiles of cells. 6) Visualisation of relative peak expression and read coverage across genes for select cell populations.

### Peak calling

The Sierra peak calling procedure, implemented as the *FindPeaks* program, requires three inputs: 1) a BAM file containing the data for the entire experimental run, such as produced by the 10x Genomics CellRanger software, 2) the reference (GTF) file used for the mapping and 3) a file containing splice junctions from the experiment (BED format). The BAM file must include the error corrected cell and UMI barcode tags. Although many peak callers have been developed to work with DNA sequencing data, e.g. ChIP-seq analysis, we found it was not appropriate for RNA sequencing in single cells due to the presence of exon junction reads that result in peaks spliced across the genome.

To make the Sierra peak caller splice aware, we first extract the splice junctions from the BAM file using ‘regtools’ [46] (≥ version 0.5.1). The advantage of this approach is that by extracting splice junctions directly from the data, we do not depend on existing transcript annotations, enabling discovery of novel splicing events. Using the set of identified junctions, we separate read coverage into “within junction” and “across junction” sub-sets and perform peak finding for each coverage sub-set separately. The “within junction” coverage comes from reads that align inside the identified junctions, i.e. not on junction coordinates (Figure S9). In most cases, the “within junction” reads will correspond to intronic genomic regions. The “across junction” coverage comes from remaining reads following removal of the within junction reads. In most, but not all, cases, these reads overlap with the exonic genomic regions.

Using the above input files, we perform the following steps (illustrated in Figure S9) one gene at a time:

1. The read coverage of the gene is extracted.
2. The next step is to identify a list of reliable junctions. To control for spurious splicing events, we first remove junctions with less than a specified number of junction reads. By default this is set to either 50 reads, or 5% of the maximum coverage for the gene, whichever is highest. These values are defined in *FindPeaks* using the *min.jcutoff* and *min.jcutoff.prop* parameters, respectively. The use of two criteria and stringent thresholds for filtering junctions means Sierra is relatively robust to splicing noise in genes with varying levels of read coverage.
3. To find peaks in the “across junction” regions, the following steps are applied:

a. Remove junction regions and piece together remaining coverage.
b. To call a peak, we find the genomic locus in which the local read coverage is maximum and fit a Gaussian to the read count (per base pair) using the R *nls* (nonlinear least squares) program, parameterised with the following function: *y* = *k* × *exp*(−1/2 × (*x* − *mu*)^2^*/sigma*^2^), where *y* is read coverage, *k* is peak maximum, *x* is position, *mu* is the centre of the peak location and *sigma* controls the width of the peak. We assume peaks are approximately 600 bp in width and fit a 600 bp region in each iteration. We initialise *mu* to be 300 (denoting the peak location in the center of the provided region), *sigma* to be 100 and *k* to be the maximum peak coverage. We consider a peak called if the *nls* function returns a fit without error. The called peak location is set to range from three standard deviations upstream and to three standard deviations downstream from the centre of the peak, defined as *s* − 300 + *mu*, where *s* is the site of maximum coverage for that region. Alternatively, Sierra supports Gaussian curve fitting using the maximum likelihood method, implemented using the R *mle* function. We compared peaks called using *nls* and *mle* and found them to be highly comparable, with an average of 92% of called peaks identified by both methods across 10 datasets tested (Figure S10A). We also compared runtime for calling peaks with the *nls* and *mle* method, but found no obvious speed benefit to either method (Figure S10B). In this study we used the *nls* method for Gaussian curve fitting.
c. The coverage is set to zero at the location of the fitted peak to allow the next site of maximum coverage to be identified. Steps 3b and 3c are iterated until one of two criteria have been met: either 1) a specified proportion of the read coverage has been assigned to called peaks for the gene, or 2) the maximum peak coverage is below some threshold. To evaluate the first criteria, the *FindPeaks* function takes in two parameters, *min.cov.prop* (default 5%) and *min.cov.cutoff* (default 500), which define the proportion of read coverage and total coverage, respectively. The first criteria is met when both of these thresholds are passed. The second stopping criteria relates to the maximum size of the peak. Again, two thresholds are required to be met: an absolute threshold (defined by the parameter *min.peak.cuff*; default 200) and a relative threshold (*min.peak.prop*; default 5%) defining the ratio of the current peak height relative to the maximum peak height for the gene.
4. To find peaks in the “within junction” regions, we repeat the procedure described in steps 3b and 3c for each of the “within junction” regions.

### Peak merging

Peaks are defined by their position in the genome and peaks called from multiple independent datasets are merged into a unified set of peak coordinates prior to UMI counting. The presence of peaks called across splice junctions presents a specific challenge, as some peaks may overlap but represent distinct biological signals. As an example, Figure S11A indicates the location of three peaks called from the *Cxcl12* gene across two datasets. While ‘peak 3’ is clearly distinct, peak 1 spans multiple exons, and contains peak 2. Despite the overlapping coordinates, peak 1 and peak 2 represent the expression of alternative transcript isoforms. As a result, not all overlapping peaks should be merged. Instead, the following procedure is applied to generate a unified set of peaks.

For a given gene, we evaluate the distance between any two peaks across datasets by calculating the absolute difference between start and end coordinates divided by the width of the peak. We convert the distance to a “similarity score” between 0 and 1 by taking one minus the distance, setting any negative values to 0. A score of 1 indicates a direct correspondence between start/end coordinates and 0 indicates that the difference between start and end coordinates is greater than the width of the peak. This calculation is illustrated in Figure S11A.

In order to determine a similarity score threshold for merging peaks, we calculated similarity scores between peaks from the GFP^+^ experiment (which contains two sequencing datasets) and TIP (3 datasets). For each peak, we plotted the similarity score for its best match from the comparison dataset. We found for all comparisons that the similarity scores formed a bimodal distribution (Figure S11B-E), such that on average 95% of peaks could either be matched with a similarity score ≥ 0.75, or considered clearly not matching as indicated by a score of 0. Given the drop-off in similarity distributions observed < 0.75, a default threshold of ≥ 0.75 is used to match peaks.

This process is run for all pairwise combinations of datasets to merge. That is, for each pair of datasets, peaks are compared both ways and two peaks are considered matched if at least one has a similarity score ≥ 0.75, with a relaxed criteria for the second peak (allowing a 25% deviation from the threshold). For peaks that are matched, the union of the start and end coordinates are taken to create a final merged peak.

### UMI counting

To perform UMI counting in single cells, Sierra requires a set of unified peak coordinates from *FindPeaks* or *MergePeakCoordinates*, a GTF file of gene positions, a scRNAseq BAM file and a white list of cell barcodes. We extract alignments for each gene using the GenomicAlignments package (≥ version 1.14.2), then count the overlaps between the peak coordinates and the alignments using the *countOverlaps* function in GenomicRanges [47] (≥ version 1.30.3). Extra filtering is performed to ensure only cells that are in the barcodes white list are counted. The final peak to cell matrix is output in matrix market format.

### Detecting differential transcript usage

We test for differential transcript usage using the differential exon usage testing method DEXSeq [25] (≥ version 1.24.4), which was originally developed for bulk RNA-seq. We adapted DEXSeq to scRNA-seq after transforming the single-cells to be tested into a small number of pseudo-bulk samples. Instead of testing for differential exon usage of genes between groups we use DEXSeq to test for differential peak usage within genes between groups. The use of pseudo-bulk samples allows for computational efficiency of testing. Given two sets of cells to be compared, we first build some *n* number of pseudo-bulk profiles for each of the cell sets by randomly assigning cells into *n* groups and summing their peak counts. By default, the value of *n* is set to 6 (see below). We use the *DEXSeqDataSet* function in DEXSeq to build a DEXSeq object, where *countData* is the raw peak counts, *groupID* is set to gene names, and *featureID* is a set of unique gene-peak numbers. We run the *estimateSizeFactors* function, setting the *locfunc* option to use the *shorth* function from the *genefilter* R package [48], as it is suggested in the DEXSeq documentation that the *shorth* function may provide better results for low counts. After estimating size factors we follow the standard DEXSeq pipeline for testing differential usage. The function to perform DU testing is implemented in Sierra as *DUTest*, and contains options to filter for genomic feature types and peaks annotated as near A-rich regions prior to DU testing.

The main parameter that is introduced here is the *n* value determining the number of pseudo-bulk profiles. In order to decide on a default value, we considered the impact of different values of *n* when running DU testing on the TIP and PBMC 7k datasets by re-running tests for 10 different seeds (and therefore different random assignments of cells to pseudo-bulk profiles) for three values of *n*: 3, 6 and 10. We evaluated the number of DTU genes obtained (Figure S12A,B), the consistency of those results across the ten seeds (Figure S12C,D) and finally computational time taken for a DTU test (Figure S12E,F). We found that, on average, there was a small increase in the number of DU genes detected, as well as the consistency, with an increased value of *n*; however, the computational time increased drastically when increasing *n* from 6 to 10. Thus, a value of *6* maintains a fast computational time while returning a similar number and consistency of DTU genes to a higher *n* value of 10, and was therefore selected as the default value for Sierra.

### Peak annotation

All reported peaks were annotated to identify overlapping feature information within gene transcripts using the Sierra function *AnnotatePeaksFromGTF*. This function takes peak coordinates (GRanges format) and identifies if the boundaries are within a UTR, exon or intron of known genes (supplied as a GTF formatted file). In cases where a gene has multiple transcripts, an annotation hierarchy is applied such that when peaks overlap multiple features UTR > exon > intron. In cases where a peak overlaps multiple features within a single transcript all features are returned. This same function also assesses genomic sequence downstream from peaks for the existence of poly A motifs (i.e. AAUAAA), or A-rich region (defined as 13 consecutive A with up to 1 mismatch). Similarly genomic sequence upstream of peaks were assessed for the existence of a T-rich region (13 consecutive T with up to 1 mismatch) as previously identified by [28].

### Coverage plots

Cell type BAM files were extracted from the original total single cell population BAM alignment files using the Sierra function *SplitBam*. For bulk RNA-Seq data sets the BAM files had previously been imported into SeqMonk using default RNA-seq settings. Coverage information was extracted from SeqMonk files by exporting wig-like files (via selecting running window generator on specific gene lists). Single-cell split BAM files and bulk RNA-Seq coverage data files were passed to the Sierra *PlotCoverage* function.

### Detecting 3’UTR shortening

In order to evaluate 3’UTR shortening, we first calculate DU peaks, selecting for peaks falling on 3’UTRs and filtering out peaks annotated as proximal to an A-rich region to enrich for real polyA sites. This is performed in Sierra using the *DUTest* function with *feature.type = “UTR3”* and *filter.pA.stretch = TRUE*. To ensure that our comparative analysis was performed on the same 3’UTR, we selected exon IDs for DU 3’UTR peaks using the GenomicFeatures [47] (≥ version 1.30.3) *threeUTRsByTranscript* function. For each DU 3’UTR peak, we compared the DU peak to the remaining peaks expressed on the same UTR. The peaks were ordered according to their proximity to the start of the 3’UTR and assigned a score between 0 and 1, with the most proximal peak scored 0 and the most distal scored 1. The ordering scores were compared between up- and downregulated peaks using Wilcoxon rank-sum test to evaluate shifts toward more proximal or distal peak expression.

### Plotting relative peak expression

For situations where overall gene expression may mask relative differences in peak usage, Sierra provides functionality for plotting the relative expression of peaks within a gene, whereby the peak expression is transformed according to the relative usage of the peaks among cell populations. For every gene *g*, which has *n* peaks (in which *n* ≥ 2), the relative expression of any peak expression value *x* within cluster *c* is computed as:

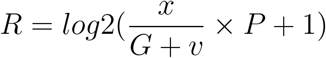

in which *x* is the original peak expression value, *G* is the mean gene-level expression of cluster *c*, *v* is a pseudo-count (a value of 1 is used in this manuscript) and *P* is the relative usage of the peak, where *P >* 1 indicates higher relative usage and *P* < 1 indicates lower usage. *G* is defined as:

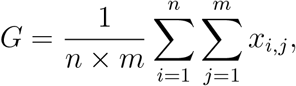

where *x_i,j_* is the expression for peak *i* and cell *j*. *P* is defined as:

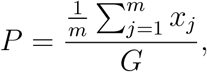

where *m* is the number of cells as above and *x_j_* is the peak expression of cell *j*.The primary motivation of this approach is to scale the individual peak expression levels based on the mean expression level each that peak and gene, within a cluster. This approach highlights the variation of peak expression within a cluster.

Finally, the relative peak expression can be visualised using similar methods for visualising gene expression, including plots on 1) t-SNE coordinates, 2) UMAP coordinates, 3) box plots and 4) violin plots. Plots are generated using ggplot2 [49] (≥ version 3.1.1).

### Clustering analyses

Gene-level clustering of the 4k and 7k PBMC datasets was performed using the Seurat R package (version 3.0.2). We applied the following quality control metrics: for both datasets, cells with > 10% UMIs mapping to mitochondrial genes were filtered out. We visualised the distribution of expressed genes and UMIs and filtered out cells with outliers. For 7k PBMCs, we filtered out cells with over 15,000 UMIs and 3000 genes and for 4k PBMCs we filtered for 10,000 UMIs and 2000 genes. For both datasets, the UMIs were then normalized to counts-per-ten-thousand, log-transformed and top 2000 variable genes were selected using the *FindVariableFeatures* program. The variable genes were used for principal component (PC) analysis, with the top 20 PCs input to the *FindNeighbors* program. A range of resolutions (*res* parameter) were applied to the *FindClusters* program with a resolution of 0.6 chosen for both the 7k and 4k PBMCs for the DTU analysis presented in this manuscript.

For the comparisons between gene-level and peak-level clustering, we used three datasets: the PBMC 7k, *Pdgfra*-GFP^+^ and TIP. For the GFP^+^ and TIP datasets, which were originally clustered in Seurat version 2, we reclustered in version 3 for the purpose of comparisons, though retaining the same PCs used for the original clustering [26]. For all datasets, the pipeline for clustering on peaks remained identical to that for the relevant gene-level clustering, such as the same number of PCs used for clustering; however, we experimented with increasing the number of highly variable features included for PC analysis, due the peaks in effect splitting genes into multiple features. For the more heterogeneous TIP data we selected 3500 features, for 7k PBMCs we used 2500 and 2000 were retained for GFP^+^. We compared the returned clusters across 3 *res* parameters: 0.6, 0.8 and 1.0. To compare gene- and peak-level clusters we used the *adjustedRand* function from the clues R package [50] (version 0.5.9).

### Mice

Mice were bred and housed in the BioCORE facility of the Victor Chang Cardiac Institute. Rooms were temperature and light/dark cycle controlled. Standard food was provided ad libitum. Mouse lines carrying targeted alleles used in this study are as follows:

1. Wild type [Inbred C57BL/6J]
2. Pdgfra^+/GFP^ [B6.129S4-Pdgfratm11(EGFP)Sor/J]
3. RaMCM [B6.Cg-Pdgfra¡tm1.1(cre/Esr1*)Nshk]
4. tdTomato [B6.Cg-Gt(ROSA)26Sortm9(CAG-tdTomato)Hze/J]

Mice carrying tamoxifen-inducible Cre recombinase (MerCreMer), knocked-in Pdgfra locus (RaMcM^+/-^) were crossed with tdTomato^+/+^ mice to generate Pdgfra lineage-tracer mice (Pdgfra^tdTom^). Male mice aged 8-12 weeks were used for the experiments.

### Tamoxifen treatment

Reporter activation in Pdgfra^tdTom^ mice was induced by 3 intraperitoneal injections of tamoxifen on consecutive days at 100 mg/kg.

### Surgically induced myocardial infarction

MI surgeries were performed 1 week after the tamoxifen treatment. To induce acute MI, mice were anaesthetized by intraperitoneal injection of a combination of ketamine (100 mg/kg) and xylazine (20 mg/kg), and intubated. Hearts were exposed via a left intercostal incision followed by ligation of the left anterior descending coronary artery. Sham operated mice underwent surgical incision without ligation. Hearts were harvested for cryo-embedding or FACS sorting at 3 days post-surgery, as indicated in Results.

### FACS

Cardiac fibroblasts (PDGFRα+) and endothelial cells (CD31^+^) were isolated from wild type C57Bl/6J mouse hearts and sorted using APC-conjugated anti-PDGFRα+ (eBioscience 17-1401-81) and PE-CY7-conjugated anti-CD31 (eBioscience 25-0311-82) antibodies as described previously [36]. Cardiomyocytes were isolated from wild type C57Bl/6J mice as described previously [51].

### RNA isolation and real time quantitative RT-PCR

Total RNA from isolated cells or whole heart was isolated using TRIzol Reagent (ThermoFisher Scientific 15596026) following the manufacturer’s instructions. RNA was re-suspended in nuclease free water and RNA concentration and purity determined spectrophotometrically. 1 ug RNA was used to prepare cDNA using Quantitect reverse transcription kit (Qiagen 205313) followed by real time PCR using SYBR Green (Roche Applied Science 04707516001). For altered 3’UTR usage candidates we used oligo dT for cDNA synthesis. PCR was performed on a BioRad CFX96 Real-Time Detection System using the following conditions: an initial denaturation for 5 min at 95°C, followed by 35 cycles of 10 sec denaturation at 95°C, annealing for 10 s at 60°C and extension for 15 s at 72°C. Melting curve analyses and sequencing of the amplification products were performed to verify the specificity of the amplification. The threshold cycle (Ct) was determined and the relative quantitative expression of mRNAs were calculated using method ∆∆*ct* and normalized to HPRT as an internal control. For altered 3’UTR usage candidates quantitative expression was calculated as described in Figure S13.

### Primers used in qRT-PCR

The primers used for qRT-PCR were synthesized by IDT (IDT Inc.) and are listed in Table S5

### EdU pulse-chase experiment, immunohistochemistry and confocal microscopy

For EdU (5-ethynyl-2’-deoxyuridine, Life technology) labelling experiments in vivo, animals were injected intraperitoneally (i.p.) at 200µg g^−1^ body weight at day 3 post-MI and sacrificed after 24 hr. Hearts were fixed in 4% PFA for 3 hr and washed in PBS before being incubated in 30% w/v Sucrose/PBS overnight at 4°C. Tissues were embedded in Tissue-Tek (Sakura, 4583) and frozen on dry ice. 8 µm thick transverse sections were prepared for immunohistochemistry. Sections were washed in PBS and treated with 5% BSA, 0.1% Triton X-100 for 1 hr at room temperature. EdU staining was performed with Click-iT EdU Imaging kit (Thermo Fisher Scientific, C10337) according to the manufacturer’s instructions.

## Declarations

### Availability of data and materials

Sierra is implemented as an open source R package and is available at https://github.com/VCCRI/Sierra under a GPL-3.0 licence [52]. A copy of the Sierra source code used in this manuscript is available on Synapse under Synapse ID syn21368359 [53].

The datasets analysed during this study are available as follows. The 7k human PBMC and 4k human PBMC datasets are available from 10x Genomics [54, 55]. The TIP and GFP^+^ datasets can be found on ArrayExpress under identifier E-MTAB-7376 [56]. The Tabula Muris dataset is available from the Amazon web store at [57]. Cardiac bulk RNA-seq data is accessible through the Gene Expression Omnibus under accession ID GSE95755 [58]. Processed peak count files generated in this study are available on Synapse under Synapse ID syn21368359 [59].

### Author contributions

The study was conceived by RP, DTH, AO, JWKH and KKL. The method was developed by RP, DTH and KKL with advice from AO, JWKH and RPH. Experiments were carried out by RP, DTH, KKL and VJ with advice from AO, JWKH and RPH. RP wrote the manuscript with input from DTH and KKL. All authors reviewed and edited the manuscript. All authors read and approved the final manuscript.

### Ethics approval and consent to participate

All experimental procedures were approved by the Garvan Institute/St. Vincent’s Hospital Animal Experimentation Ethics Committee (No. 19/14) and were performed in strict accordance with the National Health and Medical Research Council (NHMRC) of Australia Guidelines on Animal Experimentation.

### Consent for publication

Not applicable

## Acknowledgments

We thank Vikram Tallapragada for performing surgeries and Alexander Ward for providing technical assistance.

## Supplementary tables legends

**Table S1.** Metrics from peak calling, counting and annotation for the 19 individual data-sets analysed is this study. Shown are 1) the number of peaks, 2) number of genes, 3) median peaks per-gene, 4-7) % of peaks falling on 3’UTRs, exons, introns or 5’UTRs, 8-11) % of peaks with a poly(A) motif according to 3’UTRs, exons, introns or 5’UTRs and 12-5) % of peaks down-stream from an A-rich stretch according to 3’UTRs, exons, introns or 5’UTRs.

**Table S2.** Comparison of overlapping detected differential transcript usage genes and differentially expressed genes for different cell-type comparisons between the PBMC 7k and PBMC 4k data-sets.

**Table S3.** Output for DTU testing between cell-types from the TIP data-set.

**Table S4.**Output for DTU testing between activated/proliferating fibroblast populations and the resting populations. DTU testing based on 3’UTR peaks, after filtering out peaks tagged as proximal to an A-rich region.

**Table S5.** List of the primers used in qRT-PCR.

