## Supplementary Figures for "Sierra: discovery of differential transcript usage from polyA-captured single-cell RNA-seq data"

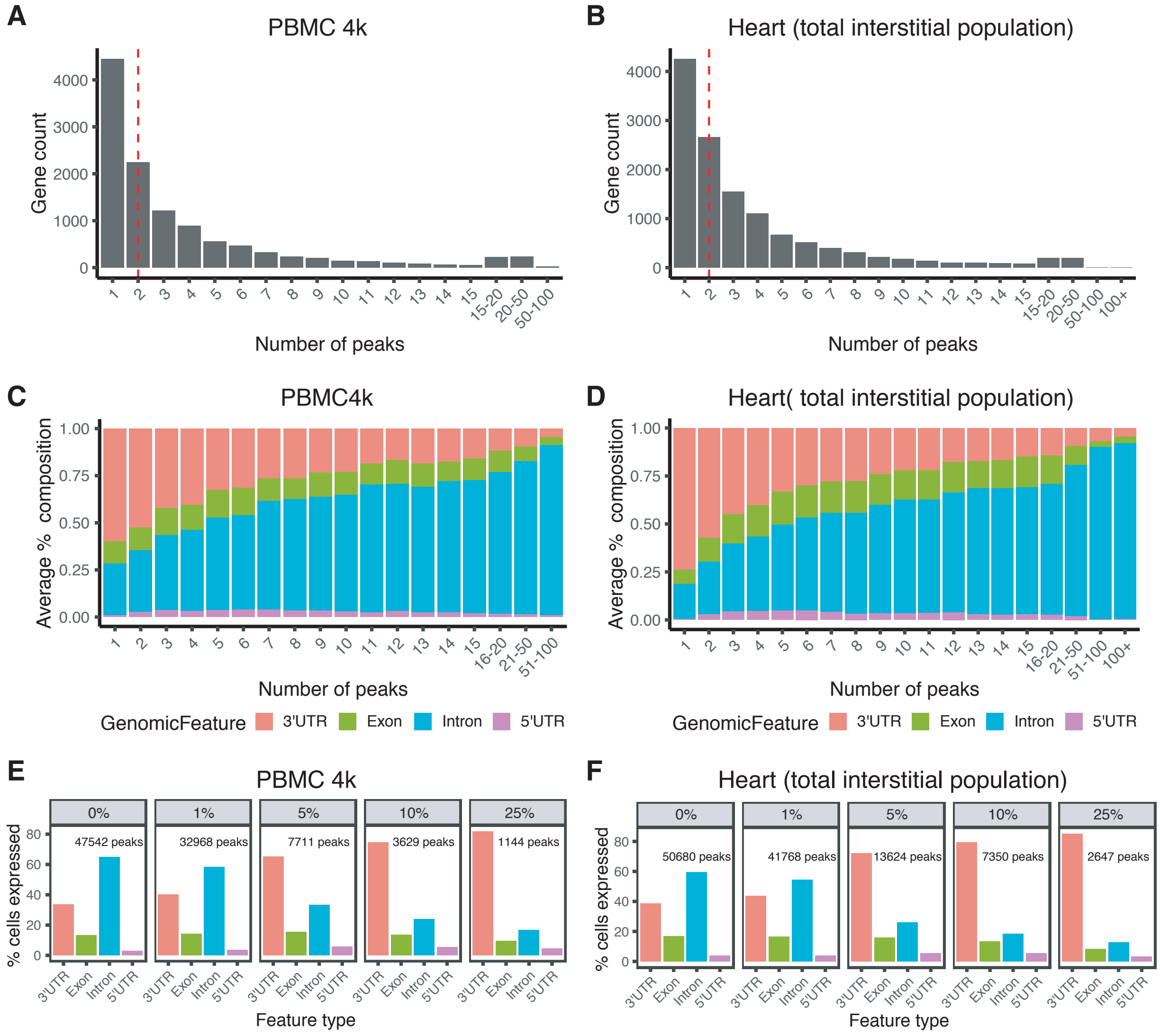


**Figure S1: Peak metrics for 4k PBMC and heart heart total interstitial population (TIP) data-sets.** (A,B) Counts of genes according to number of detected peaks where dotted red line indicates median number of peaks for the (A) PBMC 4k data-set and (B) TIP data-set. (C,D) Average composition of genomic feature types that peaks fall on, according to number of peaks per-gene for the (C) PBMC 4k data-set and (D) TIP data-set. (E,F) Percentage of cells expressing each genomic feature type with increasing stringency of cellular detection rates for peaks for the (E) PBMC 4k data-set and (F) TIP data-set.

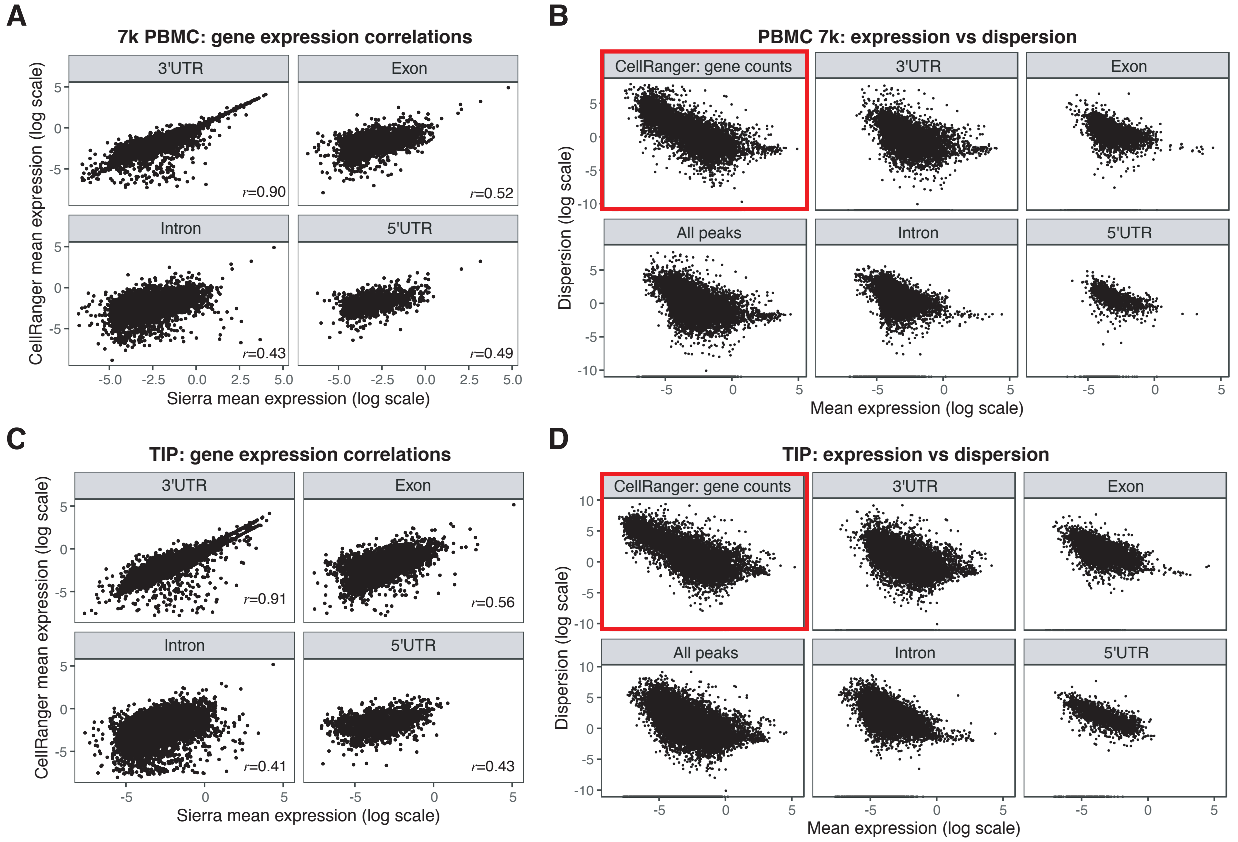


**Figure S2: Expression features of peaks comparing Sierra output to the output from the 10x Genomics CellRanger program.** (A) Correlation of Sierra-derived peak expression (summed across each gene) to CellRanger-derived gene expression for the PBMC 7k data. For both Sierra and CellRanger, the mean expression across cells is shown. For Sierra results, peak expression is stratified according to each genomic feature type – 3'UTRs, exons, introns and 5'UTRs. Pearson correlation coefficients are indicated by $r$ values. (B) Mean expression across cells compared to dispersion for the PBMC 7k data. Shown are comparisons of CellRanger gene expression (top left; red box) to Sierra peak expression (all peaks; bottom left) and peak expression stratified according to genomic feature type. (C) Expression correlation for heart TIP data. (D) Mean expression vs dispersion comparisons for heart TIP data.

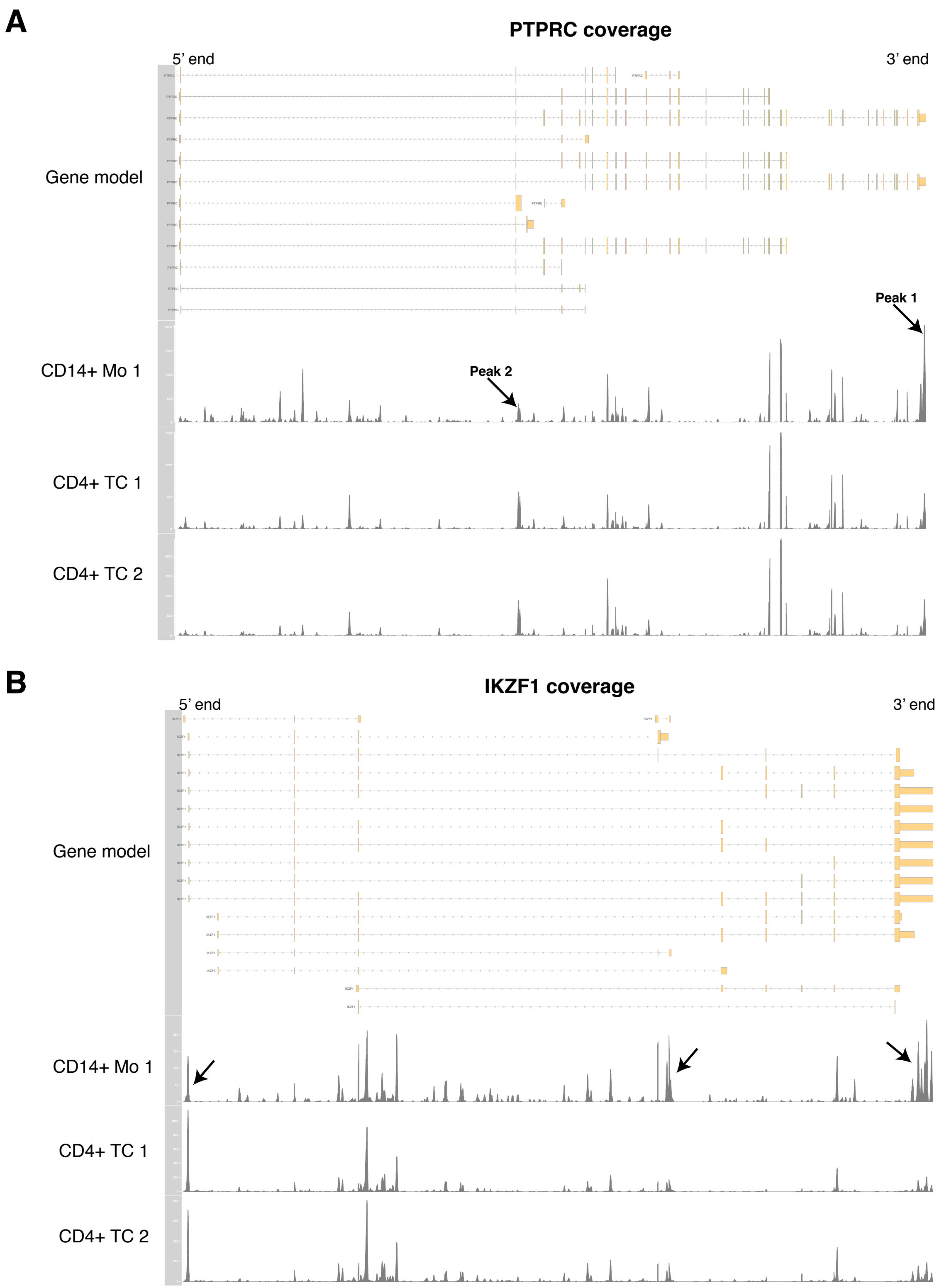


**Figure S3: Gene-level read distribution coverage plots for PBMC cell populations.** Distributions are shown for PBMC 7k populations $CD14^+$ monocyte (Mo) and $CD4^+$ T-cell (TC) populations 1 and 2 for (A) *PTPRC* and (B) *IKZF1*.

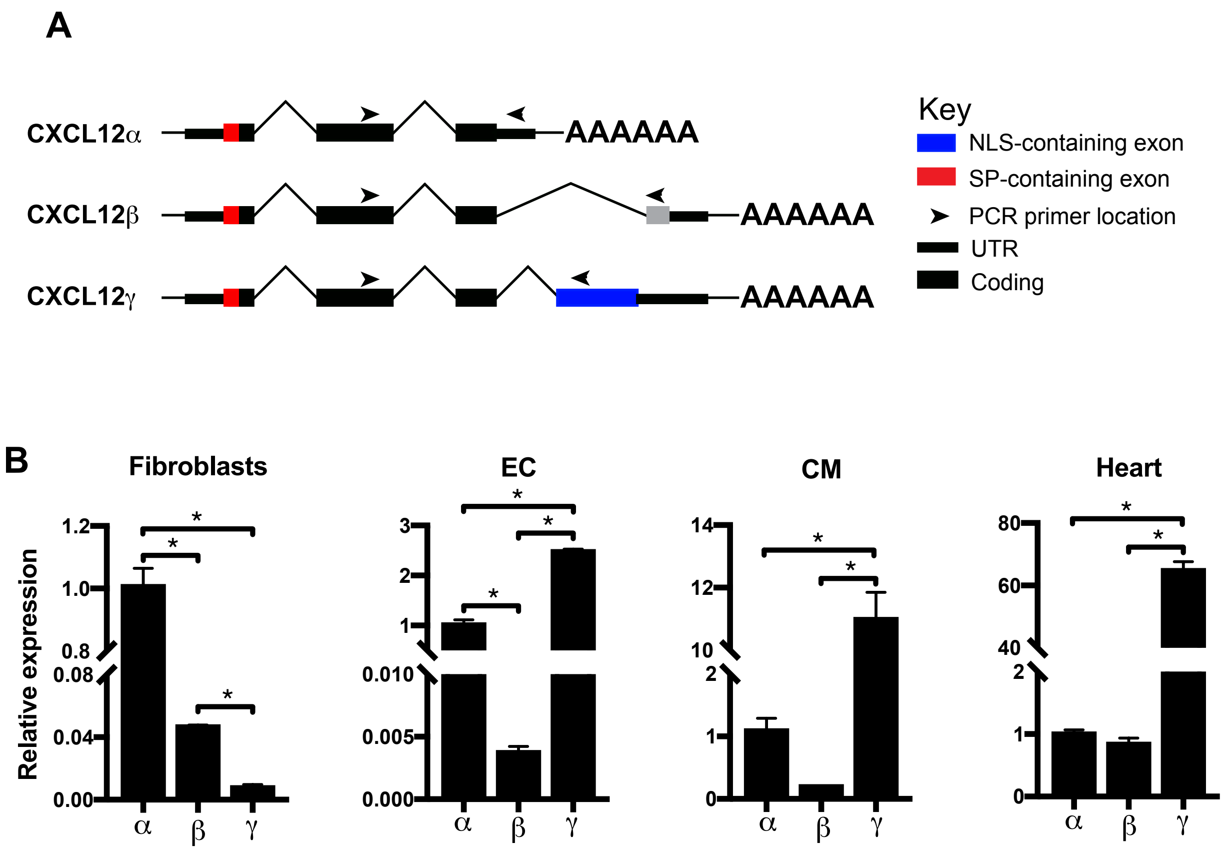


**Figure S4: Expression of *Cxcl12* isoforms in cardiac cell types.** (A) Schematic representation of isoforms of *Cxcl12* pre-mRNA showing the primer positions used for qRT-PCR. Colour code indicates the exon containing the signal peptide (SP) or nucleolus localisation signal (NLS) (B) Relative abundance of *Cxcl12* isoforms in different cardiac cell types and whole heart. Error bar shows mean values ± standard error. Stars indicate significant ($p < 0.05$) difference in expression according to a 2-tail t-test (n=3).

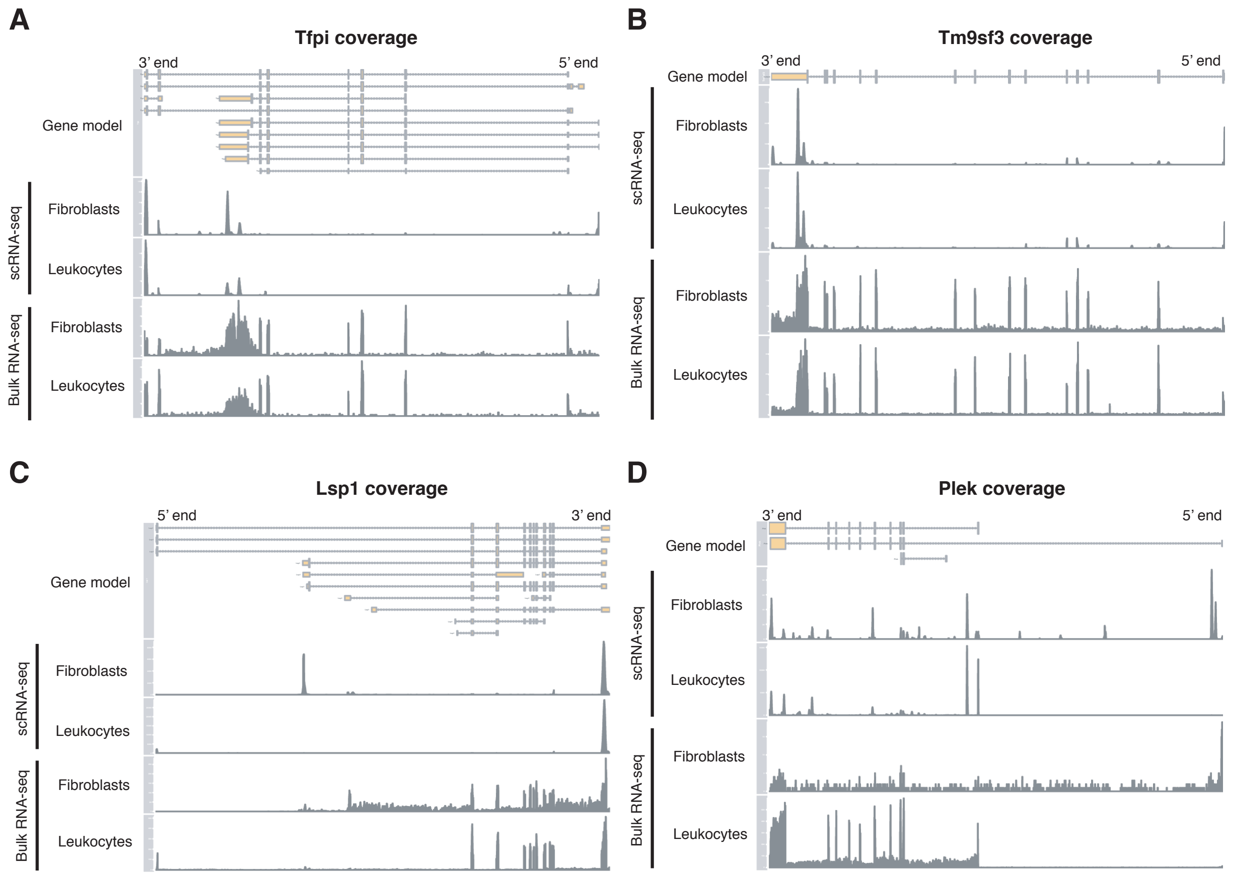


**Figure S5: Gene-level read distribution coverage plots for TIP cell populations.** Distributions are shown for scRNA-seq sham fibroblast and MI-day 3 leukocytes from TIP data compared to representative bulk RNA-seq sham fibroblast and MI-day 3 leukocyte samples for (A) Tfpi, (B) Tm9sf3, (C) Lsp1 and (D) Plek.

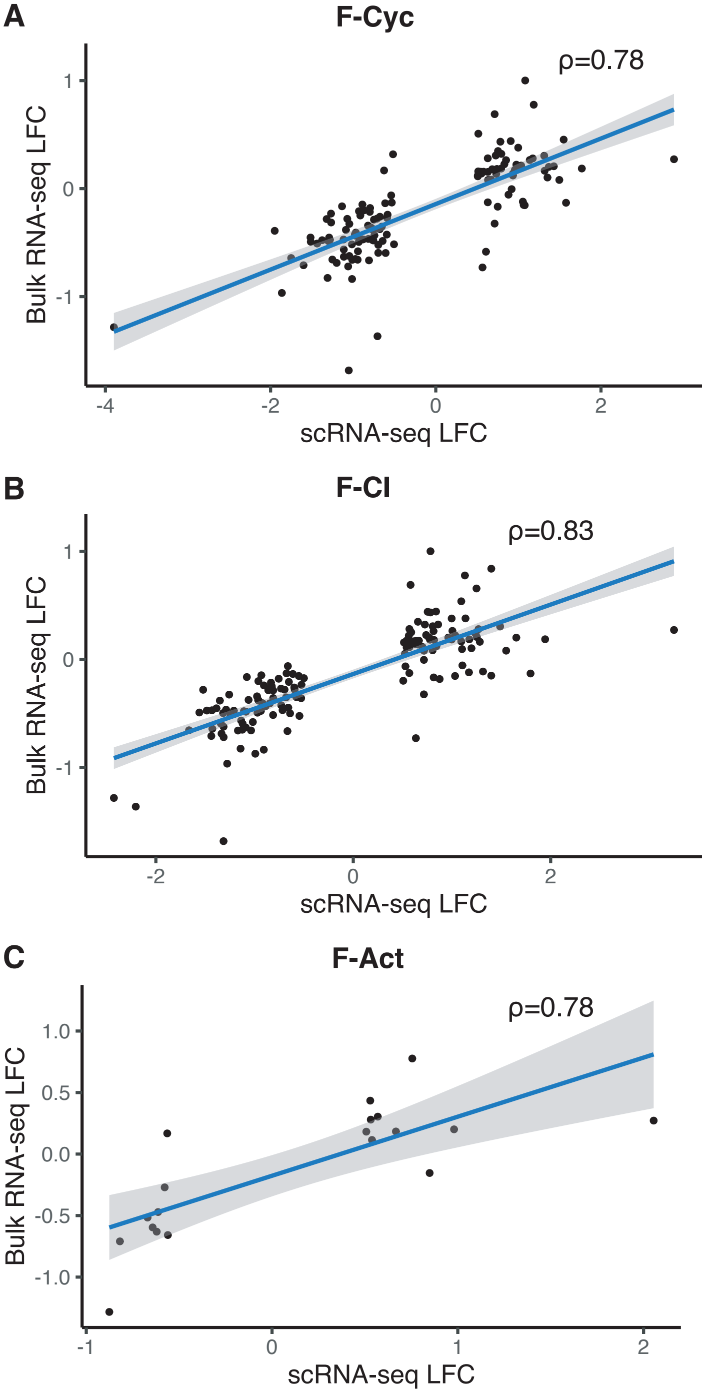


**Figure S6: scRNA-seq and bulk RNA-seq fold-change comparisons.** Log fold-change comparisons for DU peaks identified in both the bulk RNA-seq between MI and sham fibroblasts and single-cell RNA-seq for populations (A) F-Cyc, (B) F-CI and (C) F-Act compared to resting populations F-SH and F-SL. Shown is the Spearman correlation coefficient for each comparison.

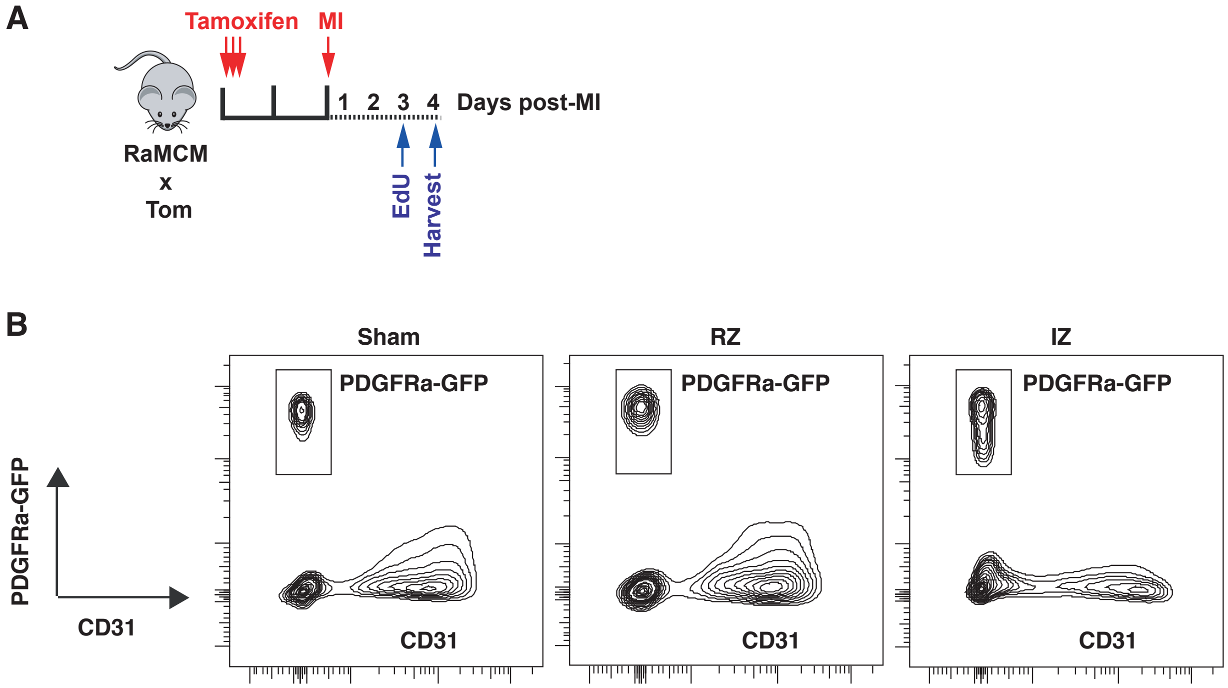


**Figure S7: EdU experimental procedure and FACS plots.** (A) Schematic showing experimental design for EdU incorporation in Pdgfra$^{tdTom}$ mice subject to MI surgery. (B) Representative FACS plots showing gated and sorted *Pdgfra*-GFP$^{+}$ CD31$^{-}$ cells from sham or indicated anatomical location of MI hearts of *Pdgfra*$^{+/GFP}$ mice – the remote zone (RZ) or infarct zone (IZ).

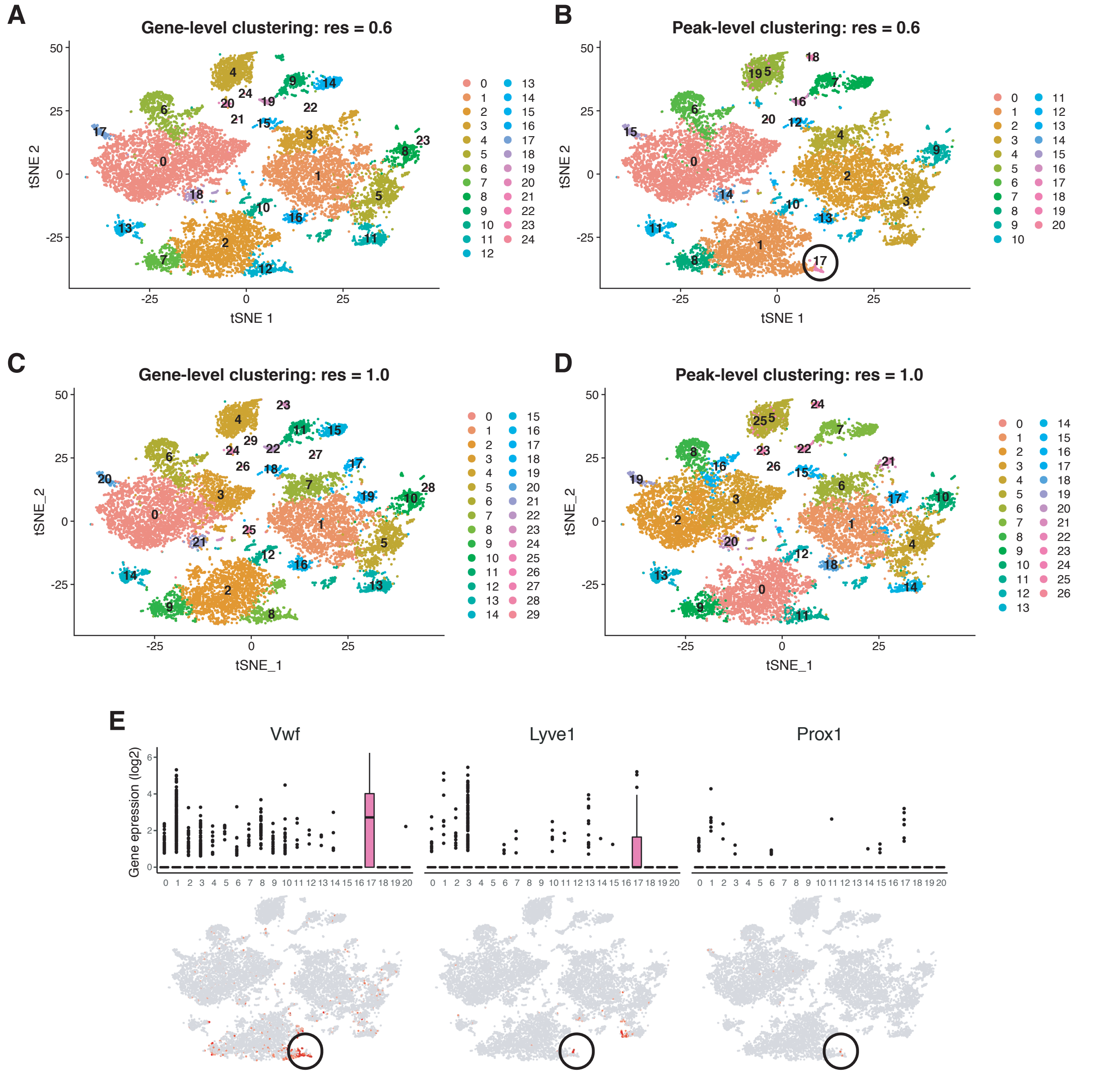


**Figure S8: Comparison of Seurat clustering between using gene and peak expression with different resolution (*res*) parameters.** (A-D) t-SNE visualisation of TIP cell clusters for (A) gene-level clustering with $res = 0.6$, (B) Peak-level clustering with $res = 0.6$, (C) gene-level clustering with $res = 1.0$ and (D) gene-level clustering with $res = 1.0$. (E) Example lymphatic EC markers upregulated in cluster '17' from peak clustering of the TIP data-set, with $res = 0.6$ .

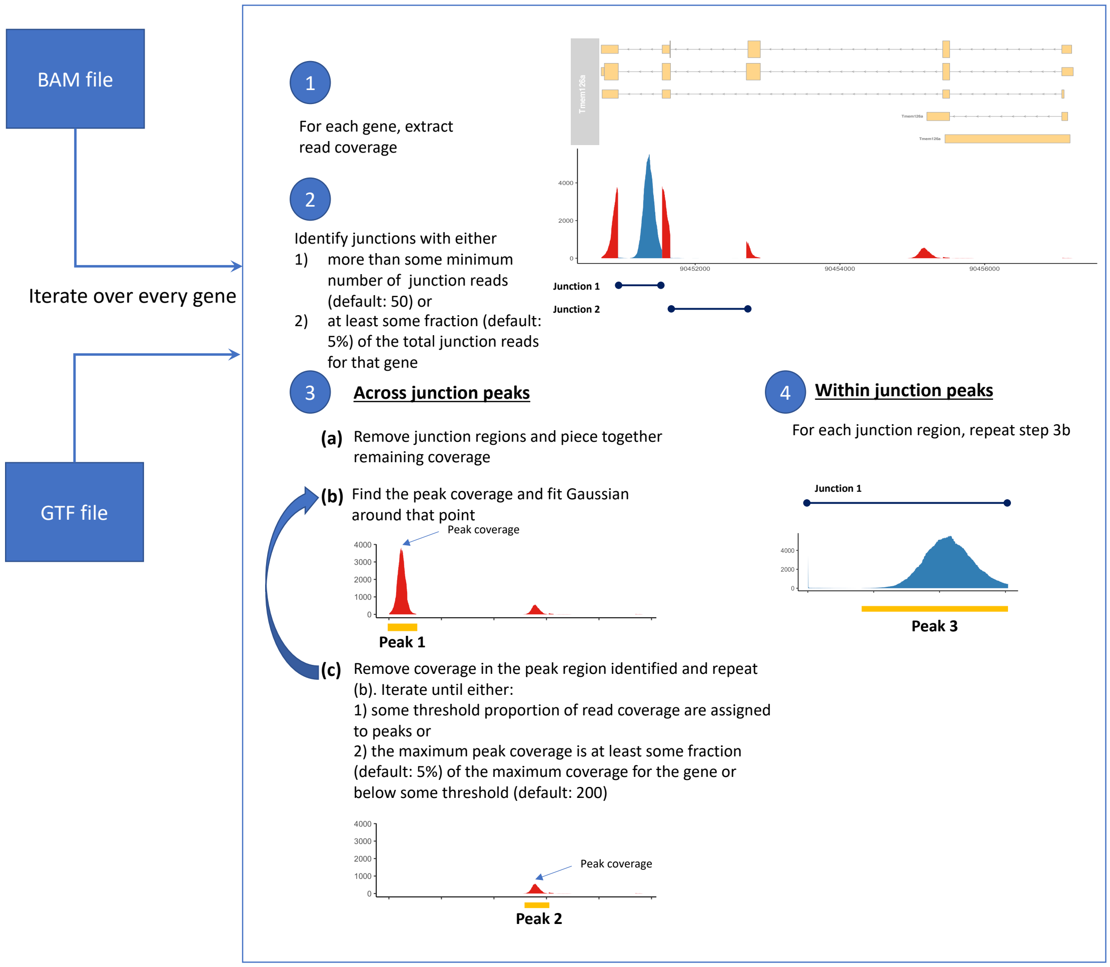


**Figure S9: Schematic for peak calling process.** A BAM file and GTF file are required inputs. Peak calling is run on each gene, according to the following steps. 1) Read coverage across the gene is extracted. 2) Junctions are identified that pass filtering thresholds. 3) Peak calling is run iteratively on "across junction" regions, removing coverage following each peak call, until a minimum amount of remaining coverage is left. 4) Peak calling is run on the "within junction" peaks up until a minimum amount of remaining coverage.

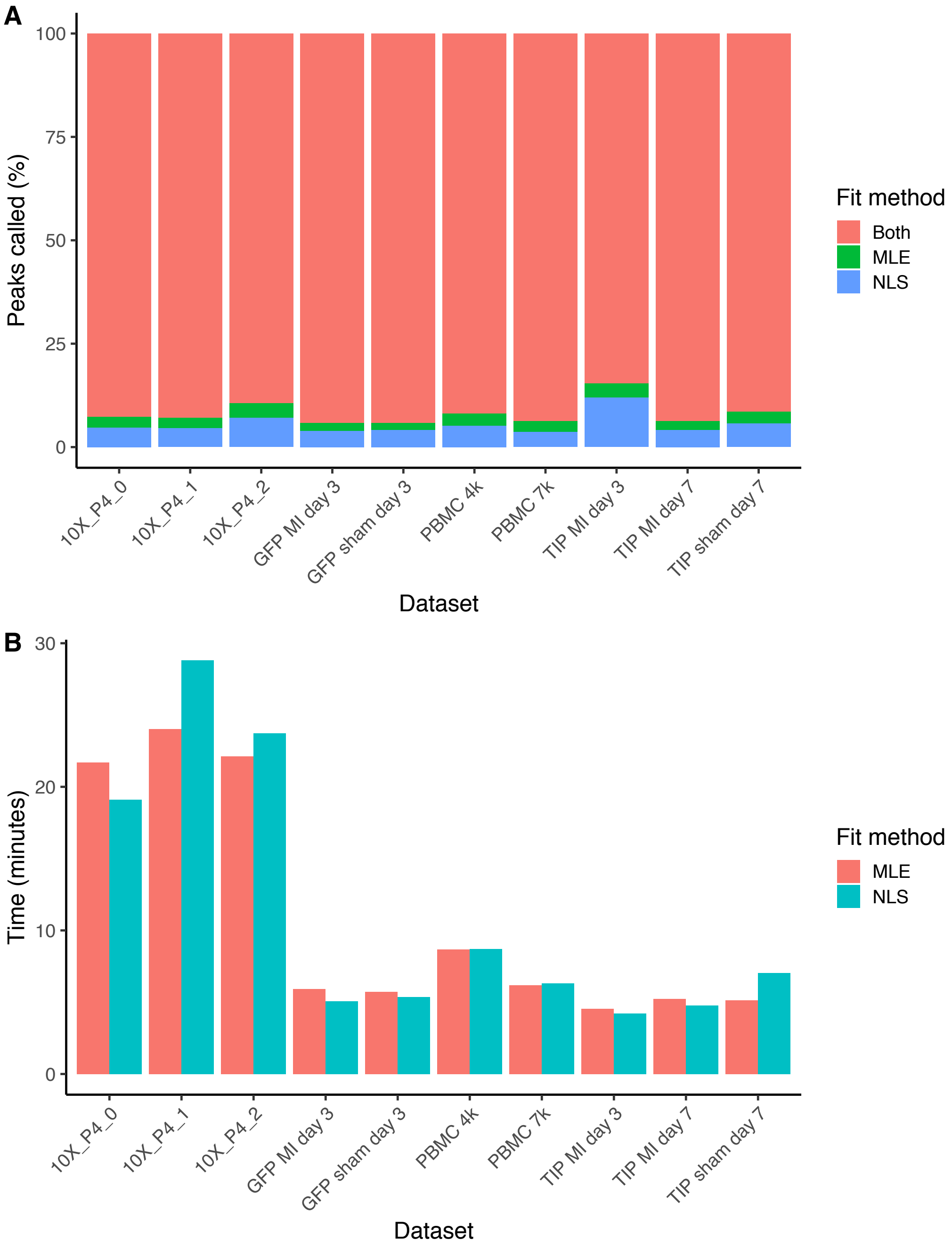


**Figure S10: Comparison of peak calling methods.** (A) Percentage of peaks called using maximum likelihood estimation (MLE) compared to nonlinear least squares (NLS). Shown are the percentage of peaks called by both methods or uniquely by either method. (B) Comparison of runtime (in minutes) for executing the peak calling procedure using either NLS or MLE.

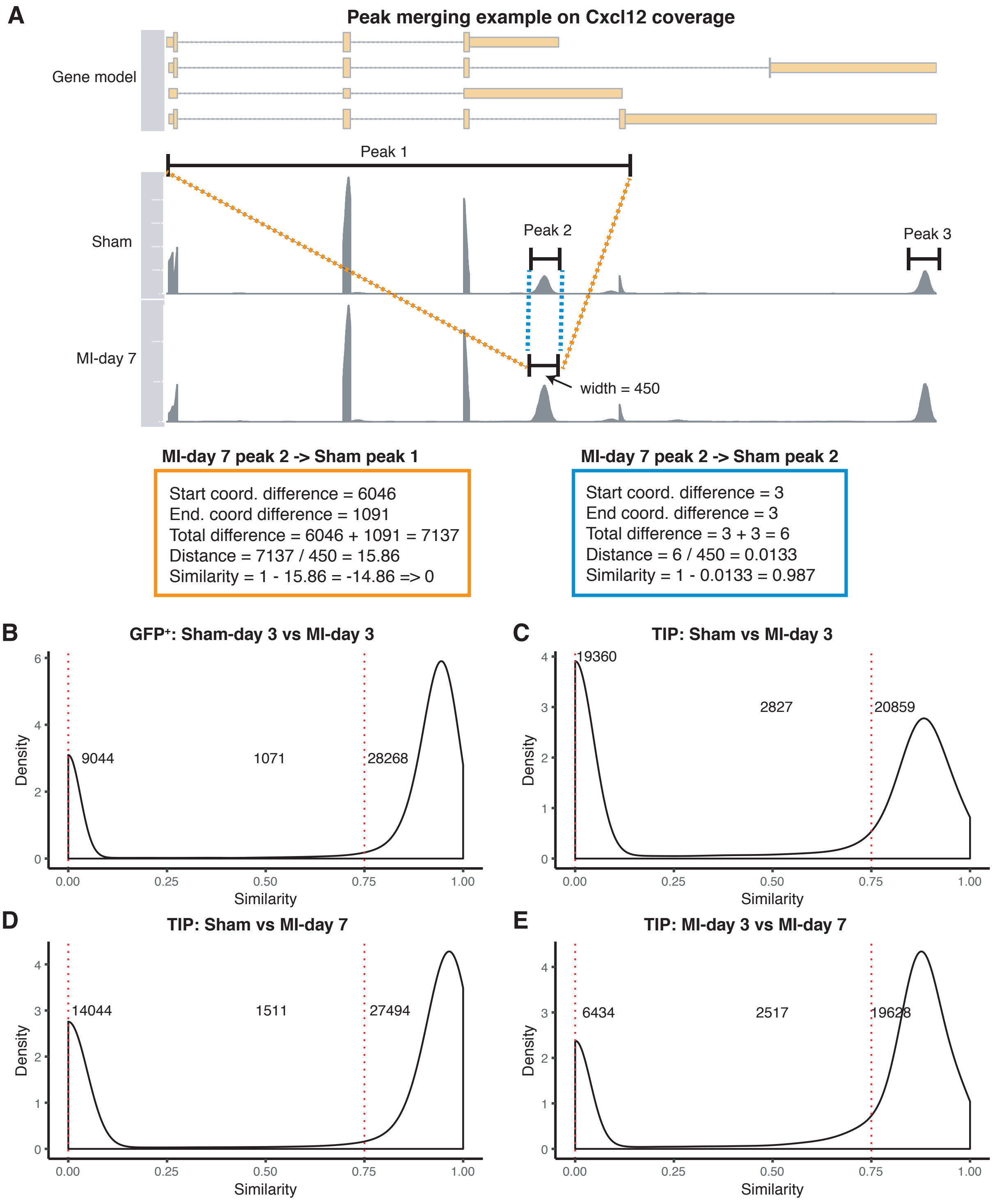


**Figure S11: Procedure and example distributions for similarity score calculations.** (A) Schematic illustrating similarity score calculation on a coverage plot of the *Cxcl12* gene for 2 data-sets: sham and MI-day 7 cardiac TIP scRNA-seq experiments. Shown are the three peaks called for *Cxcl12* in both data-sets, and example similarity score calculations comparing 'peak 2' in the MI-day 7 data-set to peak 1 and peak 2 in sham. (B-D) Distribution of peak similarity scores between pairs of data-sets. Comparisons shown for (A) sham vs MI GFP$^+$ cells, (B) the TIP sham and MI-day 3 data-sets, (C) TIP sham and MI-day 7 data-sets and (D) TIP MI-day 3 and MI-day 7 data-sets.

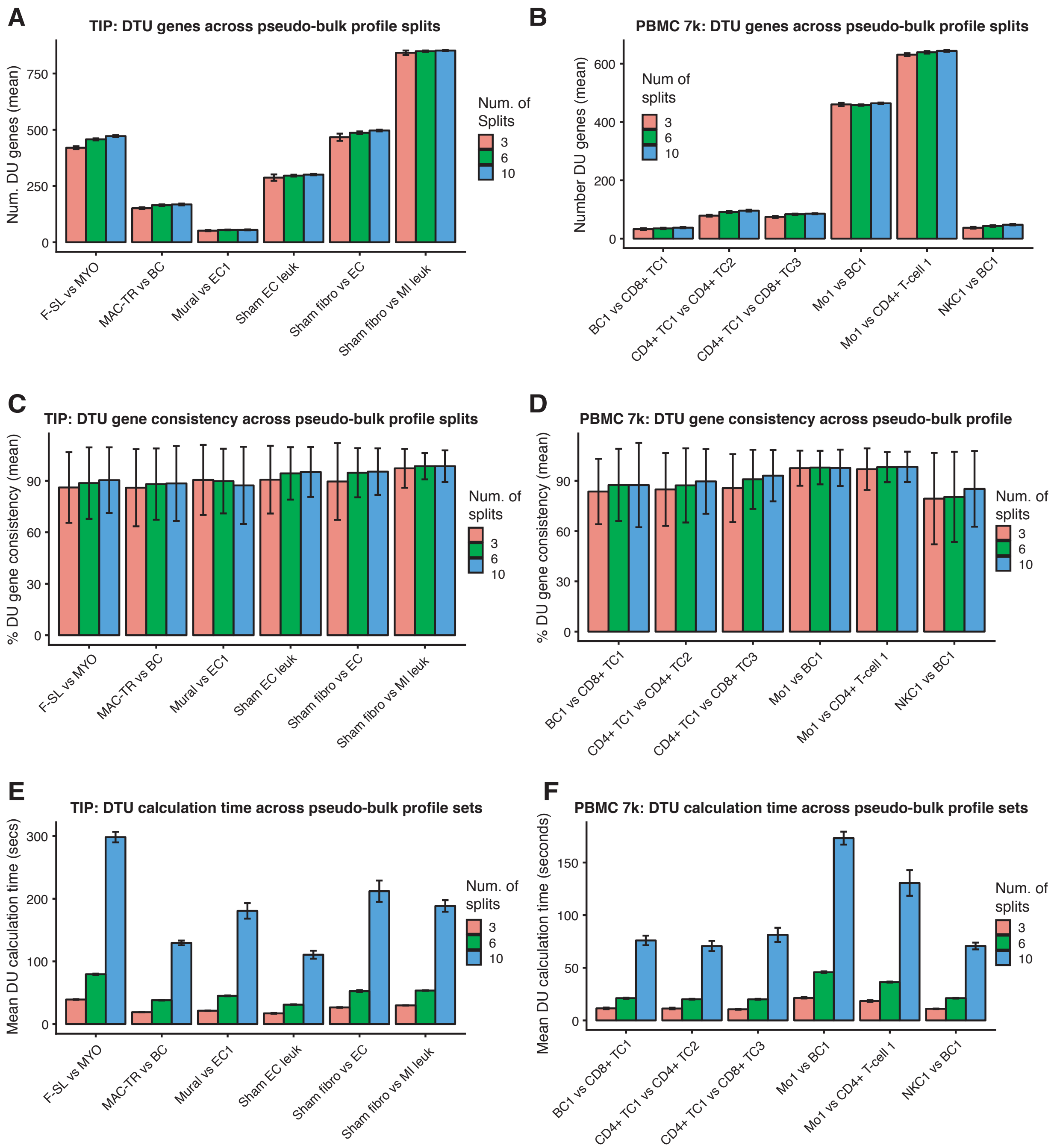


**Figure S12: Comparison of metrics from differential transcript usage testing with increasing numbers (3, 6 and 10) of pseudo-bulk profiles.** Evaluated of 6 cell type comparisons for the TIP and PBMC 7k data-sets across 10 randomised pseudo-bulk splits. Shown are mean and standard deviation for each of the following metrics: (A,B) Number of DU peaks detected per-comparison for (A) TIP and (B) 7k PBMCs. (C,D) Average consistency of DTU genes detected, where consistency is defined as the frequency of a DTU gene being detected ($p_{adj} < 0.05$) across the 10 splits for (C) TIP and (D) 7k PBMCs. (E,F) DU peak calculation time (in seconds) for (E) TIP and (F) 7k PBMCs.

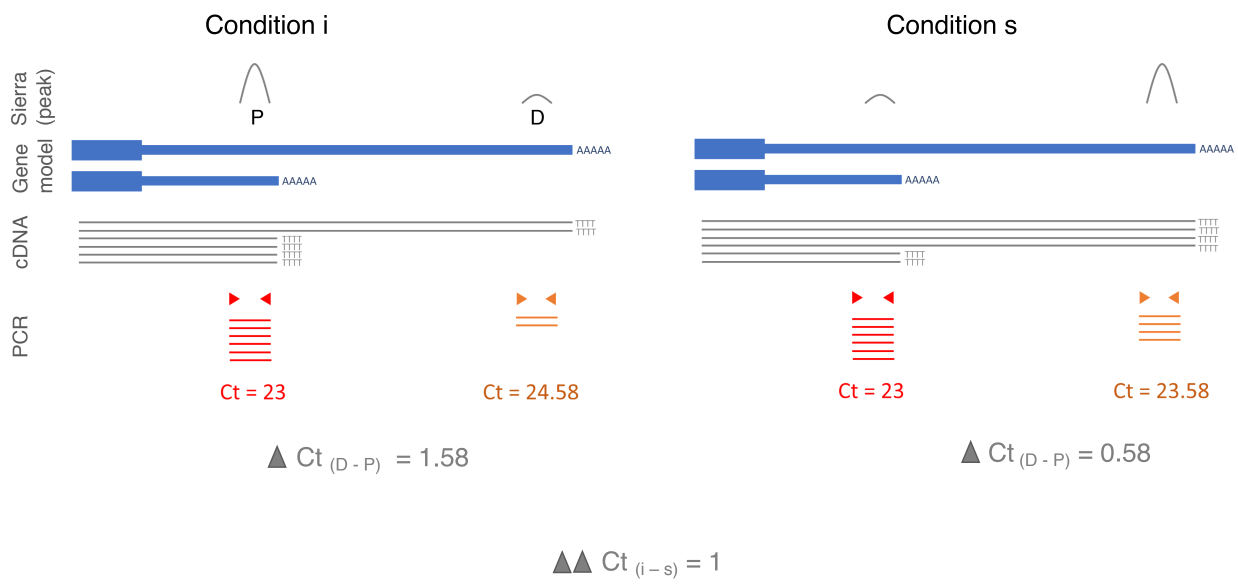


**Figure S13: Schematic of differential 3'UTR usage calculation with qRT-PCR.** Theoretical qRT-PCR example of distal vs proximal peak analysis where both condition i and condition s have the same gene expression level (i.e. 6 transcripts each. Refer cDNA which is generated from oligo dT primers). Condition i has proportionally more proximal UTR transcripts than condition s (4 vs 2 cDNA copies) and this is detected and visualised with Sierra (peak P vs peak D). PCR amplification of proximal peak will be identical for both condition i and s, but PCR amplification of distal peak will differ. Therefore when distal peak is more abundant Ct will be smaller and vice versa (refer condition i vs condition s). Delta delta Ct(i-s) is a value of one which infers that there is twice as much proximal peak (P) in condition i compared to condition s .
